# Transcriptomics supports that pleuropodia of insect embryos function in degradation of the serosal cuticle to enable hatching

**DOI:** 10.1101/584029

**Authors:** Barbora Konopová, Elisa Buchberger, Alastair Crisp

**Affiliations:** Department of Zoology, University of Cambridge, United Kingdom; Department of Developmental Biology, University of Göttingen, Germany; MRC Laboratory of Molecular Biology, Cambridge, United Kingdom

**Keywords:** insect, Orthoptera, RNA-seq, pleuropodia, embryonic organ, gland, moulting fluid, chitinase, immunity, ecdysone

## Abstract

Pleuropodia are limb-derived vesicular organs that transiently appear on the first abdominal segment of embryos from the majority of insect “orders”. They are missing in the model *Drosophila* and little is known about them. Experiments carried out on orthopteran insects eighty years ago indicated that the pleuropodia secrete a “hatching enzyme” that at the end of embryogenesis digests the serosal cuticle to enable the larva to hatch. This hypothesis contradicts the view that insect cuticle is digested by enzymes produced by the tissue that deposited it. We studied the development of the pleuropodia in embryos of the locust *Schistocerca gregaria* (Orthoptera) using transmission electron microscopy. RNA-seq was applied to generate a comprehensive embryonic reference transcriptome that was used to study genome-wide gene expression of ten stages of pleuropodia development. We show that the mature and secretion releasing pleuropodia are primarily enriched in transcripts associated with transport functions. They express genes encoding enzymes capable of digesting cuticular protein and chitin. These include the potent cuticulo-lytic Chitinase 5, whose transcript rises just before hatching. The pleuropodia are also enriched in transcripts for immunity-related enzymes, including the Toll signaling pathway, melanization cascade and lysozymes. These data provide transcriptomic evidence that the pleuropodia of orthopterans produce the “hatching enzyme”, whose important component is the Chitinase 5. They also indicate that the organs facilitate epithelial immunity and may function in embryonic immune defense. Based on their gene expression the pleuropodia appear to be an essential part of insect physiology.

## INTRODUCTION

An integral part of insect embryogenesis is the transient appearance of enigmatic glandular organs on the first abdominal segment (A1) that are called the pleuropodia (Rathke, 1844; Wheeler, 1889) (Figure 1A-C). These are paired structures that form external vesicles in some species while in others they sink down into the body wall (reviewed in e.g., Wheeler, 1889; Hussey, 1926; Roonwall, 1937). The pleuropodia are peculiarly modified limbs (Machida, 1981; Bennett, 1999; Lewis, 2000) (Figure 1D,E): their buds emerge in a line with the buds for the walking legs, but unlike the legs, the pleuropodia remain short, the majority of their cells massively enlarge and develop into a transporting-like and secretory epithelium (Bullière, 1970; Louvet, 1973; Louvet, 1975; Stay, 1977). The pleuropodia degenerate before hatching and are absent in larvae. They have been found in at least some species of nearly all insect “orders” (Figure 1F), but are absent in others, like Diptera, Hymenoptera and advanced Lepidoptera such as silkworms (e.g., Graber, 1889; Hussey, 1926; Hagan, 1931; Roonwall, 1937; Miller, 1940; Ando, 1962; Stanley and Grundmann, 1970; Ando and Haga, 1974; Bedford, 1978; Miyakawa, 1979; Machida, 1981; Norling 1982; Larink, 1983; Louvet, 1983; Kamiya and Ando, 1985; Tanaka et al., 1985; Kobayashi and Ando, 1990; Heming, 1993; Kobayashi et al., 2003; Lambiase et al., 2003; Machida et al., 2004; Rost et al., 2004; Uchifune and Machida, 2005; Tsutsumi and Machida, 2006; Mashimo et al., 2013; Fraulob et al., 2015). Perhaps because the pleuropodia are missing in the genetic model *Drosophila*, they have been neglected in recent decades. Their function has remained unclear and the genes expressed during their active stages are unknown.

**Figure 1.**
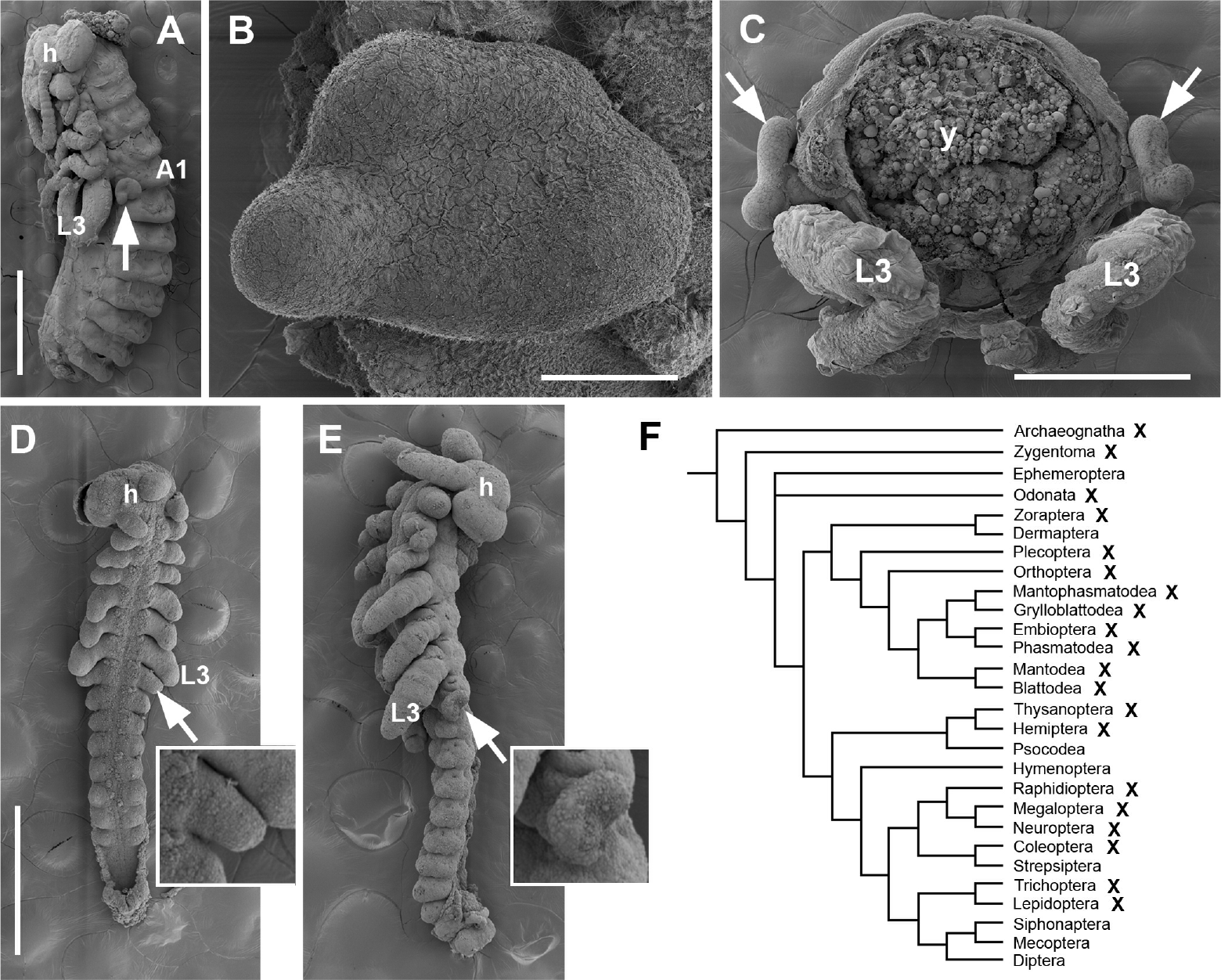
Pleuropodia are limb-derived organs on A1 of insect embryos. (A)-(C): External morphology of fully developed pleuropodia of *Schistocerca gregaria*. (A) Embryo before dorsal closure (yolk was removed). (B) Enlarged left pleuropodium. (C) Cross section through A1. (D) and (E): Pleuropodia originate by a modification of a limb bud. (D) Early embryo: all appendages are similarly looking buds. (E) Older embryo: future legs elongate and the buds on A1 start to take shape of pleuropodia. (F) Insect phylogenetic tree showing the presence of pleuropodia among “orders”. The cross marks “orders” where at least some species develop pleuropodia. Phylogeny from Kjer et al., 2016, other references in the text. (A)-(E) are scanning electron microscopy (SEM) micrographs. Pleuropodium is marked with an arrow. A1, the first abdominal segment; h, head; L3, hind (third) leg; y, yolk. Scale bars: in (A), 1 mm; in (B), 100 μm; in (C); 500 μm; in (D), for (D) and (E), 500 μm.

Eighty years ago Eleanor Slifer (Slifer, 1937; 1938) demonstrated that the pleuropodia of grasshoppers (Orthoptera) are necessary for the digestion of the serosal cuticle (SC) before hatching, to enable the larva to get out of the egg. The SC is a chitin and protein-containing sheet structurally similar to the larval or adult cuticles and is produced by the extraembryonic serosa in early embryogenesis (Goltsev et al., 2009; Jacobs et al., 2015). Shortly before hatching the inner layer of the SC (procuticle) disappears. Slifer (Slifer, 1937) showed that when the pleuropodia are removed from the embryos, the SC remains thick and the larva stays arrested in the egg. She proposed that the pleuropodia secrete the “hatching enzyme”, a substance likely similar to the cuticle degrading moulting fluid (MF) that is released by the larval epidermis under the old cuticle when the insect is preparing to moult (Reynolds and Samuels, 1996). The exact molecular composition of this “hatching enzyme” is unknown.

The endocrinologists Novak and Zambre (Novak and Zambre, 1974) argued that this would be an unusual way to digest a cuticle. During larval moulting (Nijhout, 1994) the larval epidermal cells deposit a cuticle and subsequently it is the same epidermal cells, not a special gland that secretes the cuticle degrading MF. Therefore they proposed that the SC degrading enzymes would most probably be secreted by the serosa itself. They proposed that the pleuropodia instead secrete the moulting hormone “ecdysone”, which then stimulates the serosa to secrete the “hatching enzyme”. They also suggested that the pleuropodia reach the peak of their activity in very young embryos during katatrepsis when the serosa is still present (Panfilio, 2008).

In some insects, including locusts, ultrastructural studies (Bullière, 1970; Louvet, 1973; Louvet, 1975; Rost et al., 2004; Viscuso and Sottile, 2008) have indeed shown that the pleuropodia secrete granules similar to the “ecdysial droplets” carrying the MF (Locke and Krishnan, 1973). Some of the Slifer’s experiments (Slifer, 1937) were successfully repeated by others (Jones, 1956) and a substance capable of digesting pieces of SC was even isolated from the pleuropodia (Shutts, 1952). But a proper validation by the state-of-the-art genetic methods that the pleuropodia express genes for enzymes capable to digest the SC is missing.

Here, we identified the mRNAs expressed in the pleuropodia of the locust *Schistocerca gregaria* (Orthoptera). We chose *Schistocerca* as an ideal model, because it has large embryos (eggs over 7 mm) and external pleuropodia that can easily be dissected out, and because the previous experiments testing the function of pleuropodia were carried out in orthopterans. We studied the development of the pleuropodia including using transmission electron microscopy (TEM), and by high-throughput RNA sequencing (RNA-seq) generated transcriptomes from ten morphologically defined stages. We performed differential gene expression analysis between the pleuropodia and similarly aged hind legs. For mapping of reads we assembled a transcriptome from whole embryos. The goal of this paper was to investigate whether the observed gene expression profile of the pleuropodia is consistent with the idea that these are organs for the secretion of the “hatching enzyme”. We show that during their high secretory activity the pleuropodia express genes for cuticle degrading chitinase and proteases that were previously identified in the MF. This supports the “hatching enzyme hypothesis” (Slifer, 1937; 1938).

## RESULTS

### Development of pleuropodia in the course of *Schistocerca* embryogenesis

Before we could start exploring the genes expressed in the pleuropodia of *Schistocerca* we needed to understand how these organs develop in the locust, when they are fully differentiated and show activity. Cytological study of developing pleuropodia in grasshopper embryos was previously carried out by Slifer (Slifer, 1938), but the light microscopy that she used does not provide sufficient resolution to distinguish the fine ultrastructure of the cells. Ultrastructure of pleuropodia by TEM has been described for several insects (Bullière, 1970; Louvet, 1973; Louvet, 1975; Stay, 1977; Louvet, 1983; Rost et al., 2004; Viscuso and Sottile, 2008), but a chronological study is missing for *Schistocerca* or any other orthopteran.

Under our conditions *Schistocerca* embryogenesis lasts 14.5 days (100% developmental time, DT) (Figure 2A, S1). We followed the development of the pleuropodia from the age of 4 days (27.6 % DT), when all appendages are similar looking short buds, until just before hatching, day 14 (Figures 2B, S2-S3). Simultaneously, we followed the development of the hind leg, which we used for comparison (because pleuropodia are peculiarly modified legs).

**Figure 2.**
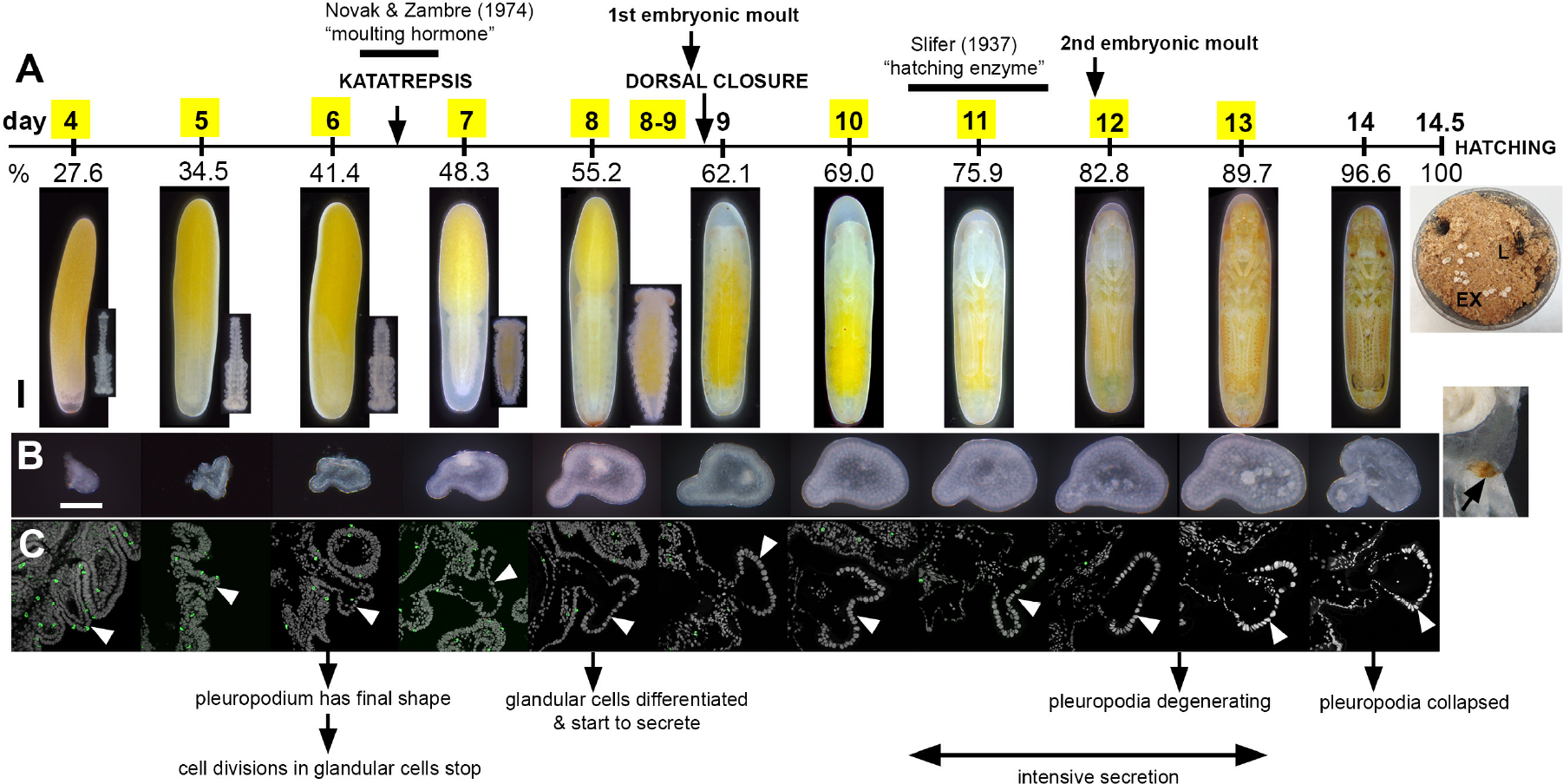
Summary of the development of pleuropodia in *Schistocerca* embryos. (A) Scheme of *Schistocerca* embryogenesis marking key developmental events in the embryos and timing of the two experiments on pleuropodia. Numbers above the scale are days from egg-laying, numbers below the scale are percent of embryonic developmental time. Yellow boxes indicate the stages that were sampled for RNA-seq. Eggs with the developing embryos at each stage are shown below the scale, insets for the 4-8 day stages show the embryo dissected out from the egg. (B) External features of the developing pleuropodia; after hatching part of the stretched exuvia is shown; the degenerated pleuropodium is marked with an arrow. (C) Paraffin sections through the pleuropodium and surrounding tissue. Pleuropodia are marked with arrowheads. PH3 (green) detects cell divisions in the immature glandular cells (tip of appendage bud) on day 4 and 5, not in later stages. The pleuropodial stalk cells, haemocytes entering the pleuropodia and cells in other tissues were labeled. Nuclei (grey) enlarge from day 6. The text below the pictures refers to the main events in the glandular cells. EX, exuvia; L, larva. Scale bars: in (A) (eggs), 1 mm; in (B), 0.2 mm. Background was cleaned in photos in A (see Materials and Methods).

We traced cell divisions in the pleuropodia by using Phosphohistone-3 as a marker (Figure 2C). The glandular cells were labeled only in the days 4 and 5. From day 6 onwards no cell divisions were detected and the nuclei started to enlarge as the cells became polyploid (Grellet, 1971). The pleuropodial stalk cells, haemocytes entering the pluropodia and cells in the other embryonic tissues kept dividing.

Although the pleuropodia get their final external mushroom-like shape just before the embryos undergo katatrepsis (day 6; 41.4% DT) (Figure 2A,B), we found by TEM (Figure 3) that the glandular cells fully differentiate only later, shortly before dorsal closure (day 8; 55.2% DT) (compare the undifferentiated cells in Figure 3F-I, with differentiated cells in Figure 3J-P). At that time these cells form a single-layered transporting-like epithelium (Berridge and Oschman, 1972) and secretion granules inside and outside the cells become visible (Figure 3A-E, J). The granules outside of the cells first appear at the base and in between the long apical microvilli (brush-border) (Figure 3E,J). The whole pleuropodium is covered with a thin embryonic cuticle (“the first embryonic cuticle”, EC1); the tips of the microvilli produce fibrous material that is a part of this cuticle (Figure 3E) (compare with similar fibers above the leg epidermis in Figure S4).

**Figure 3.**
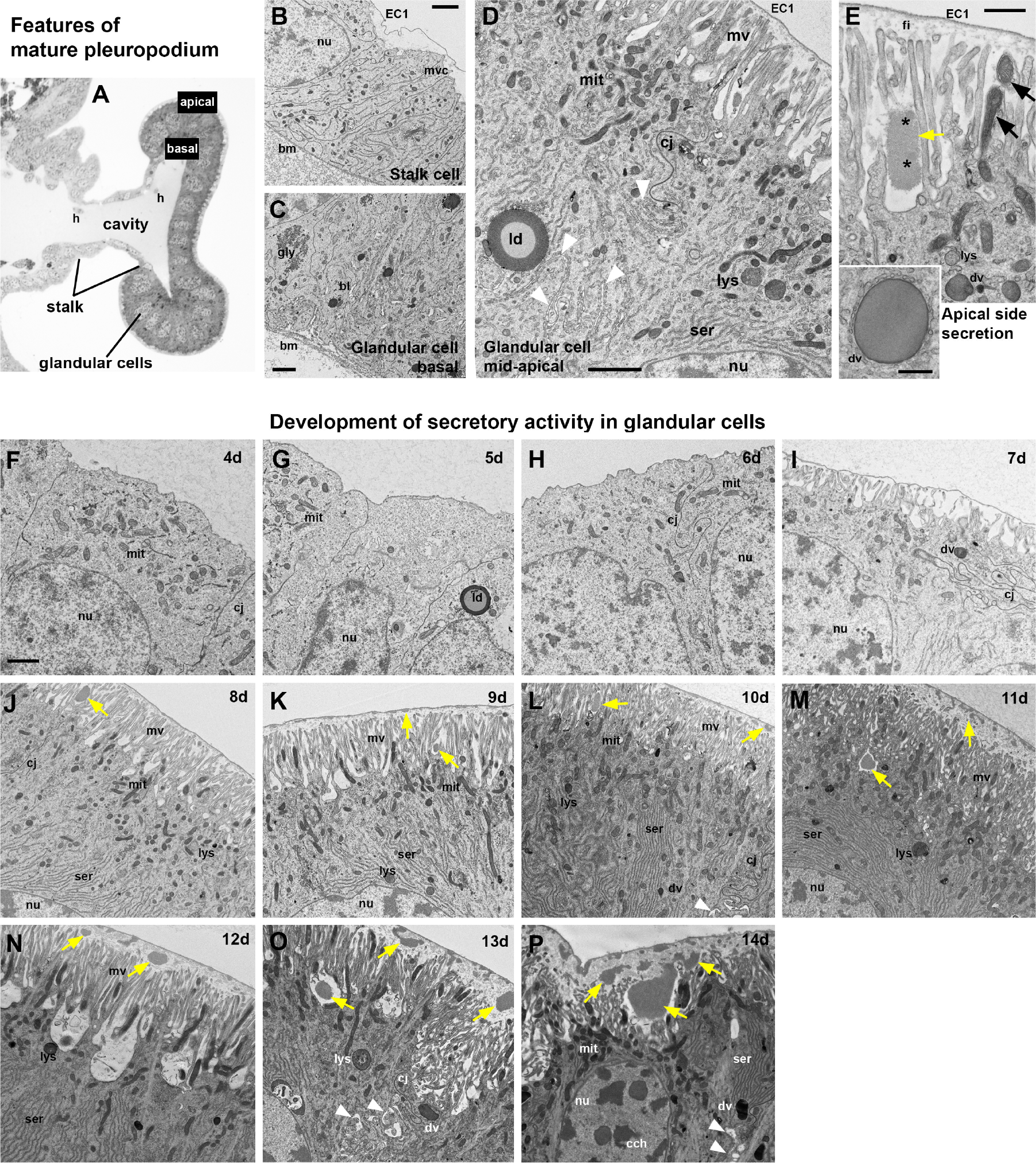
Ultrastructure of the *Schistocerca* pleuropodia. (A)-(E) Main features of the cells in the fully formed pleuropodia. Pleuropodia just before dorsal closure are shown. (A) Cross section through the pleuropodium. (B) Stalk cell. The short microvilli at the apical side are associated with the deposition of fibres in the embryonic cuticle (“the first embryonic cuticle”, EC1). (C)-(E) Glandular cells. In (D) the white arrowheads mark the spaces between neighboring cells. In (E) the black arrows mark mitochondria inside the microvilli and the asterisks mark spots of different electron-density in the secreted granule. Note that the secretion granule is located at the base of the microvilli (brush-border); the tips of the microvilli produce fibrous material that is a part of the embryonic cuticle EC1. (F)-(P) Ontogenesis of the glandular cells. Note the development of the microvilli (brush border) and the onset of secretion (appearance of secretion granules within and above the microvilli). On day 8 (J) the glandular cells are differentiated, on day 12 (N) patches of the apical side elevate, on day 13 (O) the organelles are disorganized, on day 14 (P) cytoplasm is electron dense (cells shrink), chromatin condensed, but large secretion granules are still present at the base of microvilli and above them. (A) is a toluidine blue stained semithin section, (B)-(P) TEM micrographs. Secretion granules are marked with yellow arrows. bm, basement membrane; bl, basal labyrinth (infolding of the basal plasma membrane); cj, cell junction; dv, dense vesicle; EC1, the first embryonic cuticle; gly, glycogene; ld, lipid droplet; mit, mitochondria; mv, microvilli; nu, nucleus; ser, smooth endoplasmic reticulum. Scale bars: in (B), (C), (D), (E) and (F) for (F)-(P), 2 μm; inset in (E), 500 nm.

As development progresses the secretion granules (inside and outside the cells) become more abundant and are present also above the microvilli (Figure 3K-P). On day 12 the apical side of the glandular cells changes: clusters of microvilli (usually at the borders between cells) elevate (Figure 3N). Later the cells show signs of degeneration, the chromatin condenses and the cell content becomes disorganized (Figure 3O,P). Large secretion granules are still abundant and probably released even on the last day before hatching, when the pleuropodia have shrunk and collapsed (Figures 2B, 3P).

When the embryo moults (apolyses a cuticle and secretes a new one), first at about 8.5 days and again just before 12 days (Figures 2A, S4), ecdysial droplets are present below the apolysed cuticle. These droplets are very similar at both moults (compare Figures S4F and I). They are very similar, but not identical to the granules released by the pleuropodia (Figure 4A,B). The glandular cells of the pleuropodia do not moult and keep the first embryonic cuticle (EC1) their whole life-time.

**Figure 4.**
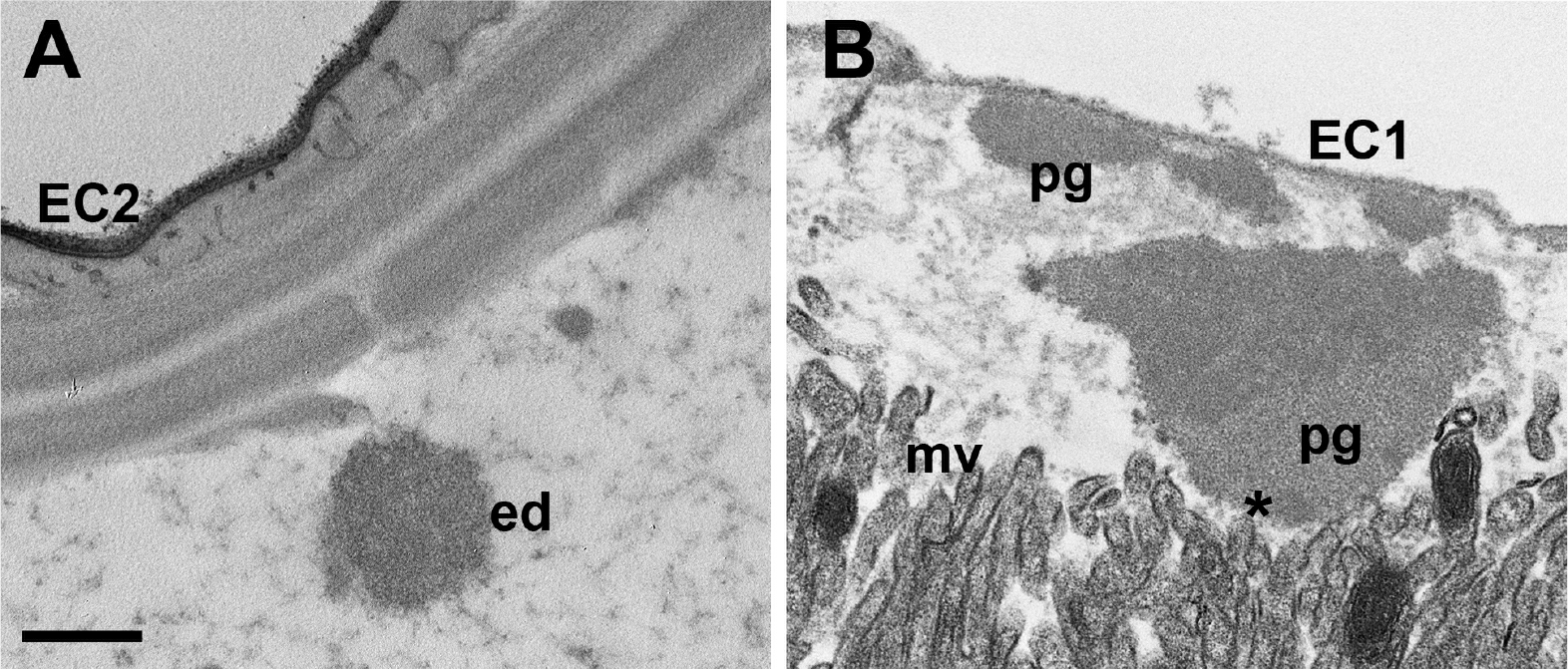
Granules secreted from the pleuropodia resemble ecdysial droplets. (A) Ecdysial droplet secreted during the second embryonic moult by hind leg epidermis. (B) Granules secreted from pleuropodia at the same developmental stage. The pleuropodial granules are typically larger, less compact and with non-homogeneous electrondensity. The “spot” of a different electron-density in the pleuropodial granules is marked with an asterisk. EC1, EC2, the first and second embryonic cuticles; ed, ecdysial droplets; mv, microvilli; pg, granules secreted from the pleuropodia. Scale bar: for (A) and (B), 500 nm.

At hatching, the larva enclosed in the (now apolysed) second embryonic cuticle (EC2) leaves the eggshell and digs through the substrate up to the surface (Bernays, 1971; Konopova and Zrzavy, 2005). Here the EC2 is shed and the degenerated pleuropodia are removed with it (Roonwall, 1937; Figure 2A).

Therefore our observations show that the timing of the high secretory activity corresponds to the stages when Slifer (Slifer, 1937) demonstrated the presence of the “hatching enzyme” (Figure 2A). Next we looked at what genes are expressed in the pleuropodia at this time.

### Generation of a comparative RNA-seq dataset from developing pleuropodia and legs of *Schistocerca*

To find out what genes are upregulated in the pleuropodia of *Schistocerca*, we applied a comparative genome wide expression analysis using RNA-seq. We generated a comprehensive embryonic transcriptome (see details in Materials and Methods) that served as reference for the analysis. This transcriptome consists of 20834 transcripts (Table S1). Its completeness was assessed using the open-source software BUSCO (version 3) (Simão et al., 2015; Waterhouse et al., 2018). 95.6%, 96.3% and 94.6% of the Metazoa, Arthropoda and Insecta orthologs, respectively, were found, a level comparable to published “complete” transcriptomes.

To gain insights into the gene expression dynamics of pleuropodia development, we dissected pleuropodia from ten embryonic stages and isolated their mRNAs. In parallel, we dissected hind legs for the same ten stages to generate a comparative transcriptomic dataset. In total we sequenced pairs of samples (pleuropodia and legs) from ten developmental stages and performed a differential expression analysis between legs and pleuropodia for each stage (Figure 2A, Table S2). A principal component analysis (PCA) confirmed that legs and pleuropodia are not only morphologically very similar at early stages, but share a common transcriptomic landscape as well (Figure 5A). The number of differentially expressed genes (DEGs) rises as development progresses (Figure 5B, Table S3).

**Figure 5.**
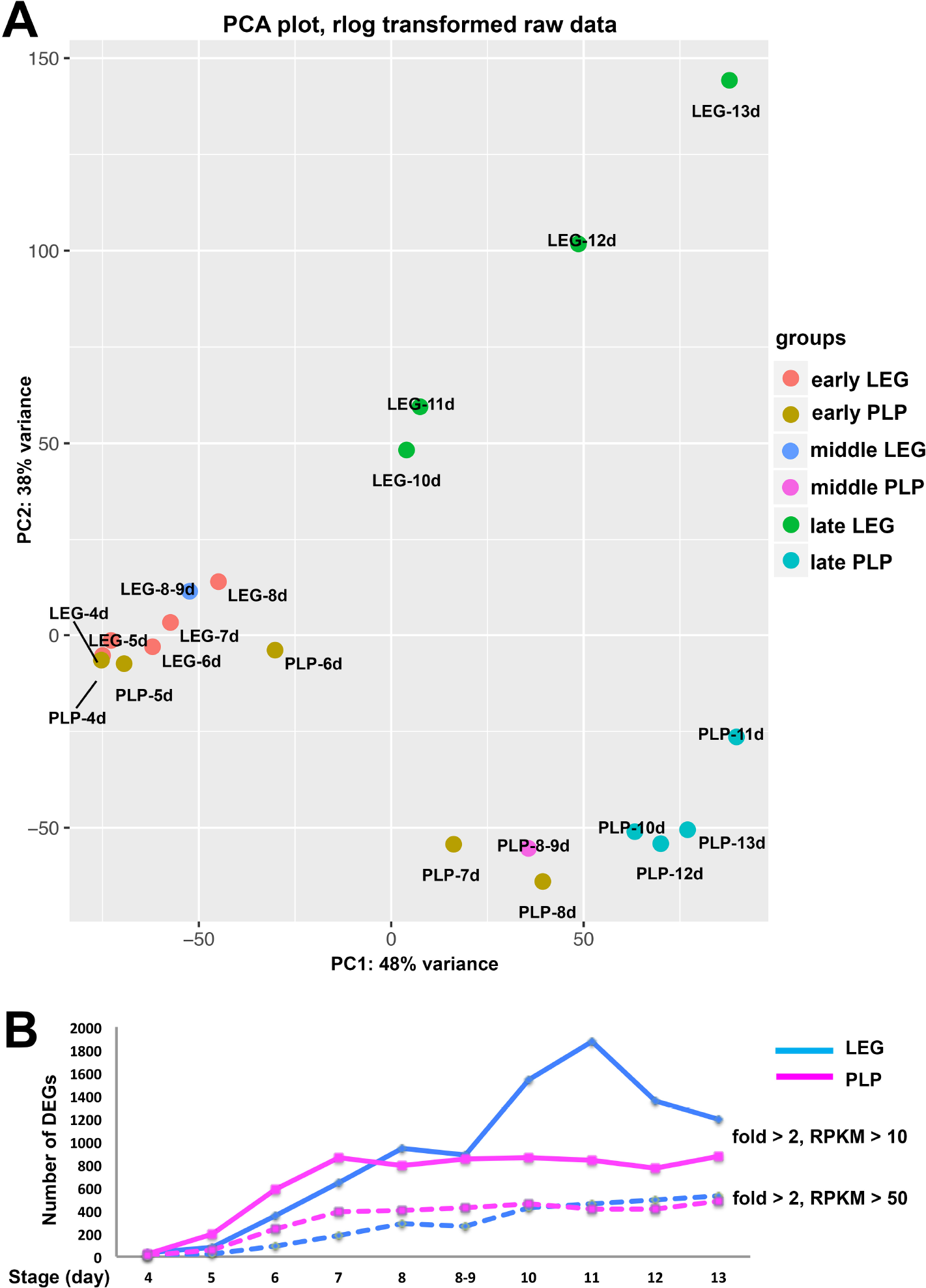
Legs and pleuropodia become genetically more different as development progresses. (A) PCA on genes expressed in legs and pleuropodia at ten embryonic stages (rlog transformed read counts). Samples from young embryos are genetically more similar and cluster together, while samples from advanced stages are genetically more distant and also separated on the plot. (B) Number of DEGs at two levels of stringency (RPKM > 10 and fold change > 2 was considered as a threshold for a gene to be differentially expressed). LEG, DEGs downregulated in pleuropodia and upregulated in legs, PLP, DEGs upregulated in pleuropodia and downregulated in legs.

For several genes whose expression dynamics in the pleuropodia were already known, such as *Ubx*, *abd-A*, *dll* and *dac* (e.g., Tear et al., 1990; Bennett et al., 1999; Prpic et al., 2001; Hughes and Kaufman, 2002; Angelini et al., 2005; Zhang et al., 2005), we confirmed that they were up- or downregulated in our RNA-seq data as predicted (Table S4). To further validate the RNA-seq dataset, we carried out real-time RT-PCR on 46 selected genes in several stages (in total in 176 cases) and got results consistent with the sequencing data (Table S5). Therefore we are confident that we can identify important factors that are relevant for pleuropodia function and development.

### Identification of genes upregulated in the intensively secreting pleuropodia

Since we wanted to focus specifically on the pleuropodia with high secretory activity we pooled the data from the samples 10, 11 and 12 days together, separately for pleuropodia and legs, and treated them as triplicates. These three samples cover the stages from the embryos after the dorsal closure, when the pleuropodia intensively release secretion granules, but are not in advanced state of degeneration (day 13) (Figures 2A, 3L-N). We performed differential expression analysis and gene ontology (GO) enrichment analysis with genes upregulated in legs and pleuropodia. We identified 781 transcripts upregulated in the pleuropodia (compared to the legs) and 1535 downregulated (Table S3). Table 1 shows the top ten percent of the most highly abundant transcripts (measured in RPKM units, “reads per kilobase of transcript per million reads mapped”) that we found upregulated in the pleuropodia.

**Table 1.** Top ten percent of the most abundant transcripts upregulated in the highly secreting pleuropodia of *Schistocerca*.

| Transcript ID | Protein | Characteristics | Immunity<sup>a</sup> | Cuticle digestion<sup>b</sup> | RPKM |  | Fold change |
| --- | --- | --- | --- | --- | --- | --- | --- |
|  |  |  |  |  | legs | pleuropodia |  |
| SgreTa0017702 | x |  |  |  | 23.07 | 15186.05 | 658.36 |
| SgreTa0007897 | C-type lysozyme | anti-bacterial protein | x |  | 42.93 | 14452.15 | 336.64 |
| SgreTa0002988 | Uncharacterized, contains DUF4773 domain |  |  |  | 15.16 | 9112.05 | 601.19 |
| SgreTa0005052 | x |  |  |  | 13.37 | 7950.98 | 594.48 |
| SgreTa0001636 | Serine protease | proteolysis | x | x | 49.38 | 7578.31 | 153.48 |
| SgreTa0008851 | Chitin binding Peritrophin-A | peritrophic matrix protein |  |  | 9.12 | 6836.42 | 749.88 |
| SgreTa0017707 | I-type lysozyme | anti-bacterial protein | x |  | 12.20 | 6712.31 | 550.26 |
| SgreTa0007042 | x |  |  |  | 7.04 | 6650.18 | 944.25 |
| SgreTa0004599 | Alpha-tocopherol transfer protein | intermembrane lipid transfer |  |  | 8.99 | 5848.12 | 650.71 |
| SgreTa0009217 | x |  |  |  | 5.03 | 5384.56 | 1070.14 |
| SgreTa0003175 | Collagen |  |  |  | 32.25 | 5220.96 | 161.87 |
| SgreTa0007886 | Alpha-N-acetylgalactosaminidase | carbohydrate catabolism |  |  | 3.85 | 4372.63 | 1134.69 |
| SgreTa0002109 | x |  |  |  | 2.20 | 3016.31 | 1372.07 |
| SgreTa0017715 | Serine protease, Snake-like | proteolysis, Toll signaling | x | x | 70.55 | 2947.46 | 41.78 |
| SgreTa0017664 | Chitinase 5 | cuticular chitin degradation |  | x | 79.32 | 2620.11 | 33.03 |
| SgreTa0002467 | Neutral endopeptidase 24.11 | proteolysis |  | x | 62.26 | 2282.01 | 36.66 |
| SgreTa0004397 | x |  |  |  | 11.21 | 2266.30 | 202.21 |
| SgreTa0002828 | x |  |  |  | 1.77 | 2188.14 | 1234.00 |
| SgreTa0006539 | Serpin, 88E-like | serine protease inhibitor | x |  | 32.42 | 2152.14 | 66.38 |
| SgreTa0001321 | Glycosyl hydrolase, Myrosinase 1-like | carbohydrate catabolism |  |  | 3.93 | 2070.40 | 527.16 |
| SgreTb0011177 | x |  |  |  | 1.38 | 1884.79 | 1369.32 |
| SgreTa0008335 | x |  |  |  | 54.24 | 1812.38 | 33.41 |
| SgreTa0003635 | Alpha-tocopherol transfer protein | intermembrane lipid transfer |  |  | 2.23 | 1800.68 | 806.99 |
| SgreTb0003860 | Serine protease, H2-like | proteolysis | x | x | 77.42 | 1727.41 | 22.31 |
| SgreTa0013418 | x |  |  |  | 0.87 | 1484.98 | 1710.66 |
| SgreTa0014009 | Angiotensin-converting enzyme | proteolysis |  | x | 65.76 | 1457.47 | 22.16 |
| SgreTa0006966 | Pro-phenol oxidase subunit 2 | immunity, melanization | x |  | 144.78 | 1347.43 | 9.31 |
| SgreTa0000425 | 6-phosphofructo-2-kinase | glycolysis |  |  | 93.52 | 1346.50 | 14.40 |
| SgreTa0003661 | Serine protease, Easter-like | proteolysis | x | x | 29.50 | 1332.79 | 45.18 |
| SgreTa0006960 | Glutamate dehydrogenase mitochondrial | nitrogen and glutamate metabolism |  |  | 172.56 | 1327.45 | 7.69 |
| SgreTa0017670 | Xaa-Pro aminopeptidase | proteolysis |  | x | 2.89 | 1322.01 | 457.96 |
| SgreTb0000759 | Cathepsin L | proteolysis, lysosomal enzyme |  | x | 105.63 | 1308.36 | 12.39 |
| SgreTa0014684 | x |  |  |  | 1.30 | 1294.87 | 994.80 |
| SgreTa0007025 | Insect pheromone-binding protein A10/OS-D | chemoreception |  |  | 1.77 | 1224.20 | 692.95 |
| SgreTa0006282 | Cytochrome P450 CYP4G102 | synthesis of hydrocarbons, anti-dehydration |  |  | 2.91 | 1196.27 | 410.93 |
| SgreTa0009515 | Sensory neuron membrane protein, 1-like | chemoreception |  |  | 3.33 | 1188.81 | 357.50 |
| SgreTa0008528 | C-type lysozyme | anti-bacterial protein | x |  | 8.61 | 1159.55 | 134.71 |
| SgreTa0009095 | Catalase | redox homeostasis | x |  | 355.15 | 1158.27 | 3.26 |
| SgreTb0039135 | x |  |  |  | 3.53 | 1119.22 | 316.71 |
| SgreTa0001486 | Lipopolysaccharide-induced tumor necrosis factor-alpha factor homolog | lysosomal degradation |  |  | 45.83 | 1109.33 | 24.20 |
| SgreTb0039012 | x |  |  |  | 14.29 | 1060.82 | 74.25 |
| SgreTa0009747 | Serpin (27-like) | serine protease inhibitor, melanization | x |  | 14.49 | 1054.67 | 72.80 |
| SgreTa0013400 | Peroxiredoxin, 5-llke | redox homeostasis | x |  | 101.10 | 1034.15 | 10.23 |
| SgreTa0017395 | x |  |  |  | 5.08 | 1004.86 | 197.64 |
| SgreTa0017712 | x |  |  |  | 15.59 | 990.41 | 63.53 |
| SgreTa0005600 | Beta-N-acetylglucosaminidase NAG2 | cuticular chitin degradation |  | x | 15.10 | 939.60 | 62.21 |
| SgreTa0000783 | Serine protease, Snake-like | proteolysis | x | x | 4.30 | 917.47 | 213.59 |
| SgreTa0006651 | Uncharacterized, contains Transcription activator MBF2 domain |  |  |  | 1.62 | 907.98 | 561.49 |
| SgreTa0017657 | Putative serine protease, K12H4.7-like / Serine carboxypeptidase | proteolysis |  | x | 2.31 | 904.26 | 391.60 |
| SgreTa0017700 | Peroxidase | redox homeostasis | x |  | 5.36 | 874.51 | 163.25 |
| SgreTa0002600 | Uncharacterized, contains DUF3421 domain |  |  |  | 0.97 | 870.73 | 894.35 |
| SgreTb0019827 | Tob | antiproliferative protein, |  |  | 141.26 | 846.86 | 5.99 |

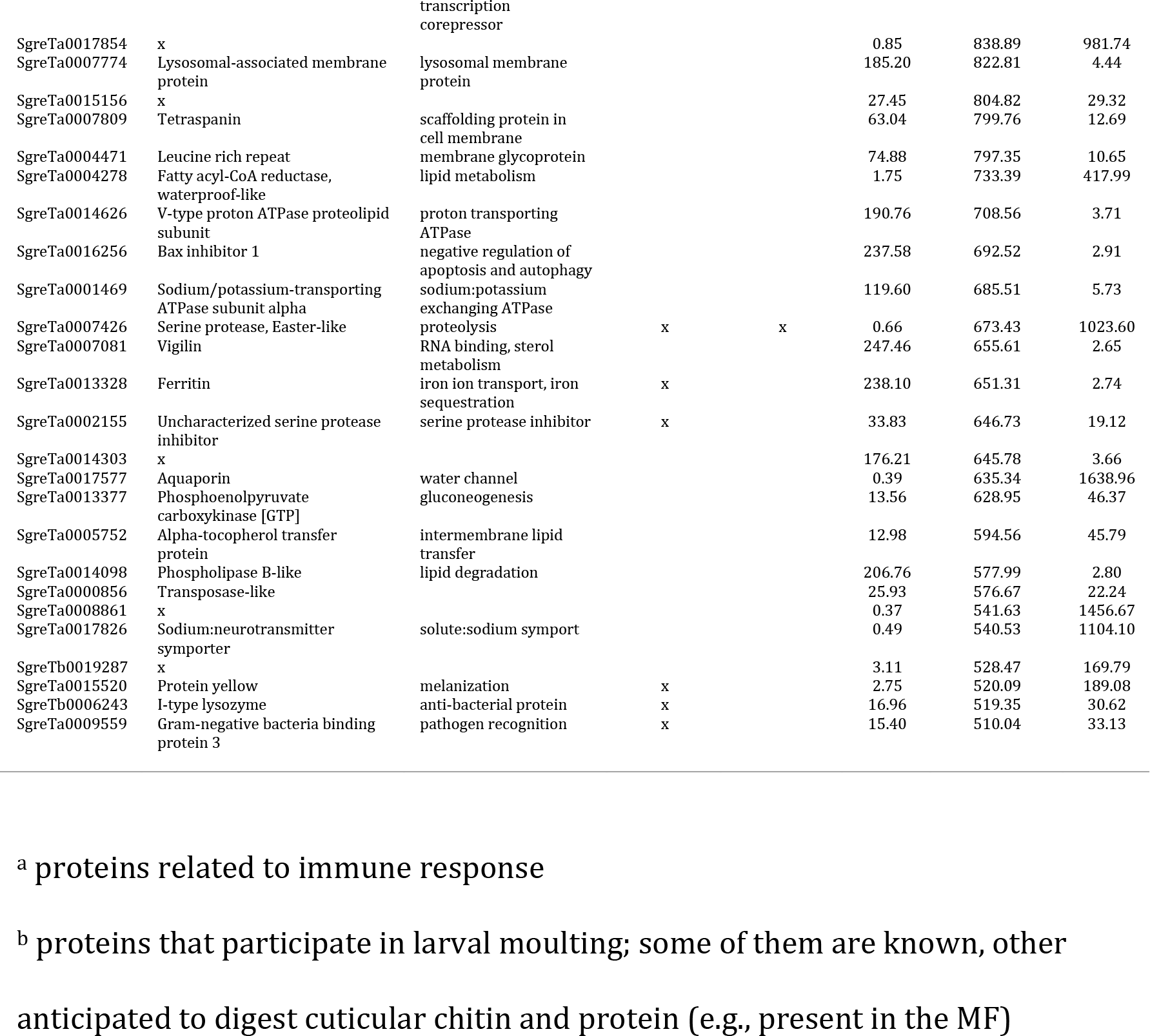

For the sake of clarity we summarized redundant GO terms in representative GO-groups (Figure 6; the full set of enriched GO terms are presented in Tables S6, S7; GOs enriched at each developmental stage separately are in Tables S8, S9). Our results show that the genes downregulated in the pleuropodia (upregulated in the legs) are enriched in GO terms associated with development and function of muscle tissue, cell division and DNA synthesis. This is in agreement with our and previous observations that the pleuropodia lack muscles, while at these stages the legs are differentiating, developing muscles and their cells are still dividing (Figure 2C). The pleuropodia downregulate genes for the development of mesoderm, which is consistent with the morphological observation that they are formed by ectodermal cells (Figure 3A).

**Figure 6.**
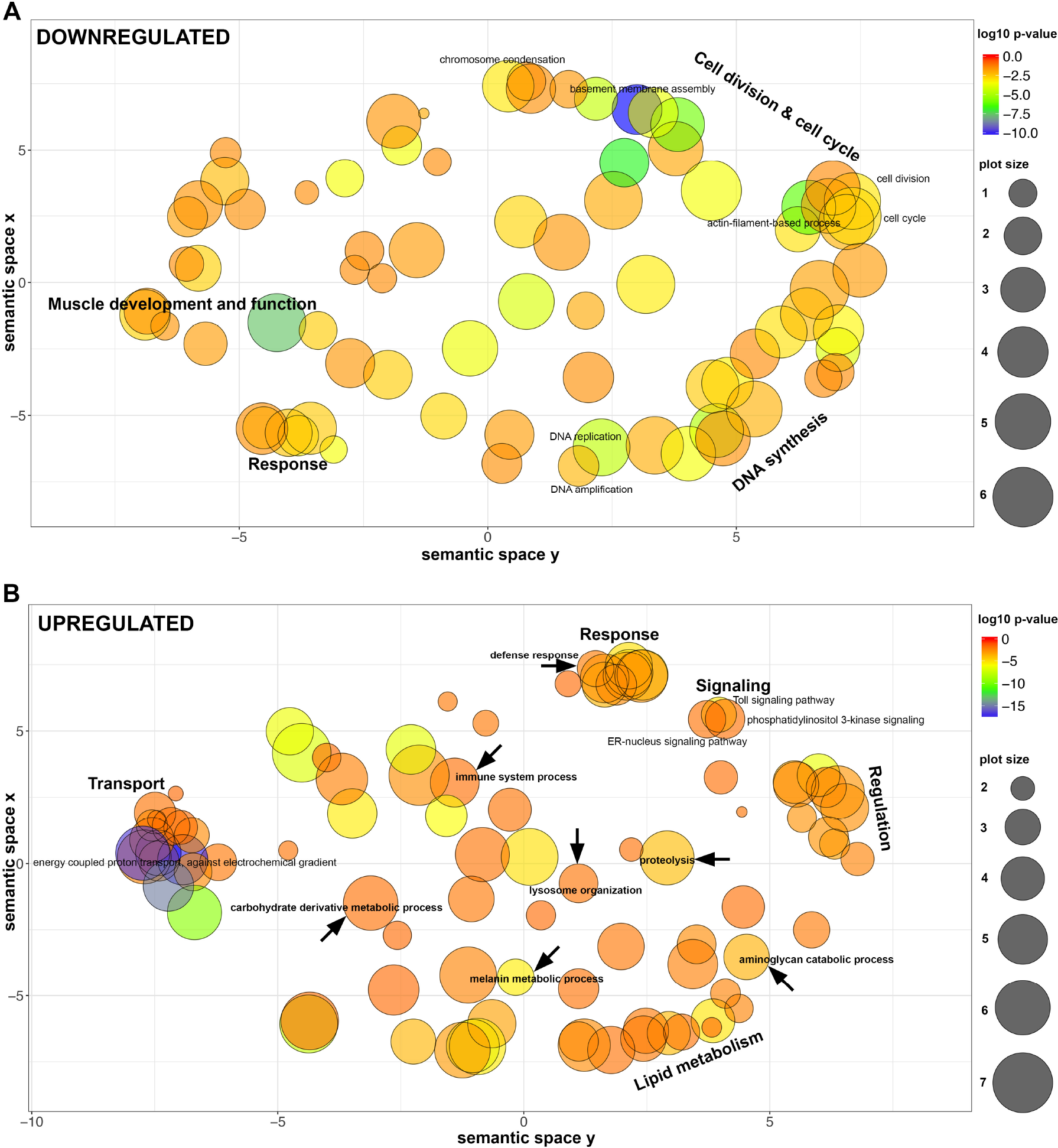
Dot plot visualization of GO terms enriched in DEGs in the highly secreting pleuropodia. Representative groups of GO terms enriched in genes that are (A) downregulated in pleuropodia (in comparison to legs) and (B) upregulated in pleuropodia. Major clusters are labeled. Relevant GOs are marked with an arrow. Bubble color indicates the p-value of the GO term, the size indicates the frequency of the GO term in the underlying Gene Ontology Annotation (GOA) database (bubbles of more general terms are larger).

The upregulated genes are primarily enriched in GO terms (Figure 6, Table S7) associated with transport, thus genetically confirming the morphological observations that the pleuropodia are transporting organs. These include genes for transporters present in typical insect transporting epithelia (Chintapalli et al., 2013), such as the energy providing V-ATPase and Na^+^, K^+^ ATPase (Table S10). We found enriched GO terms linked with lysosome organization, consistent with the observation that the pleuropodia contain numerous lysosomes (Figure 3, Louvet 1975). We also found a large cluster of GO terms associated with lipid metabolism, consistent with the abundant smooth endoplasmic reticulum in the cells. Therefore, the pool of genes expressed in the pleuropodia is in agreement with the morphology of the organs. Among the novel findings are upregulation of genes associated with immunity, as well as with carbohydrate derivative metabolism, aminoglycan catabolic process and proteolysis: these might contain genes for degradation of the SC. Next we looked at selected genes in a detail.

### Pleuropodia upregulate genes for cuticular chitin degrading enzymes

Insect cuticle is digested by a cocktail of chitin and protein degrading enzymes (Reynolds and Samuels, 1996; Zhang et al., 2014). Cuticular chitin is hydrolyzed by a two-enzyme system composed of a β-N-acetyl-hexosaminidase (NAG) and a chitinase (CHT) (Zhu et al., 2016). Both types of enzymes, a NAG and a chitinase, have to be simultaneously present for efficient hydrolysis of chitin (Fukamizo and Kramer, 1985). Previous studies have shown that only particular NAGs and CHTs are capable of efficiently digesting the type of chitin present in the insect cuticle (see below).

Insect NAGs were classified into four major classes, of which chitinolytic activity was demonstrated for group I and II (Table 2) (Hogenkamp et al., 2008; Rong et al., 2013). Our transcriptome contains four NAG transcripts, each representing one group (Table 2, Figures 7A-D, S5A, S6A). All were upregulated in the pleuropodia. Among them the *Sg-nag2* for the chitinolitic NAG group II had the highest expression (among 46 most highly “expressed” genes, Table 1) and fold change between legs and pleuropodia. The abundance of transcripts for the chitinolitic NAGs starts to rise from day 6 (Figure 7A,B) when the glandular cells in the pleuropodia begin to differentiate morphologically (Figures 1, 3). The expression profile of *Sg-nag2*, that we have chosen for validation, was similar by RNA-seq and real-time RT-PCR (compare Figure 7B and B’).

**Table 2.**
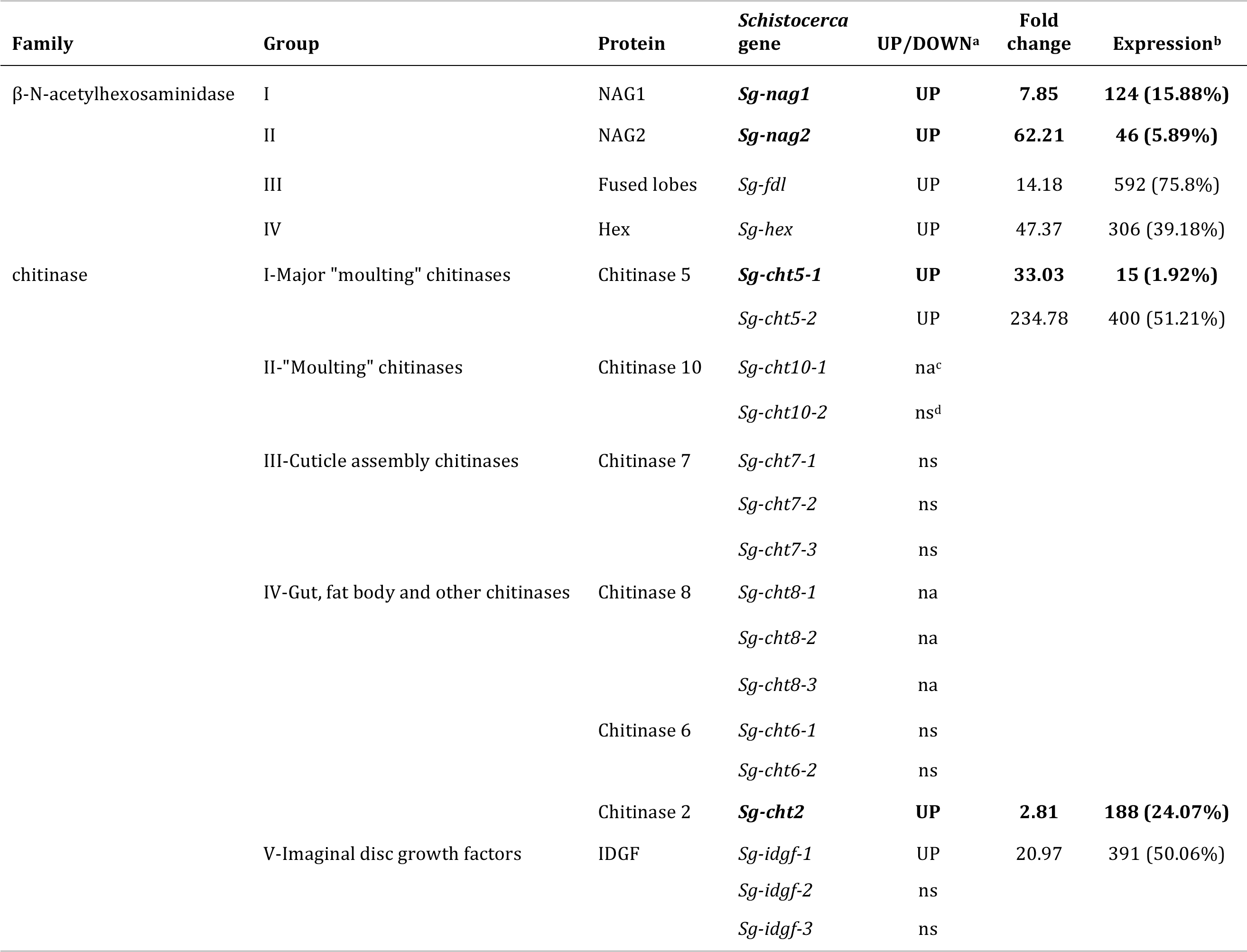
RNA-seq differential gene expression of cuticular chitin degrading enzymes in the highly secreting pleuropodia of *Schistocerca*.

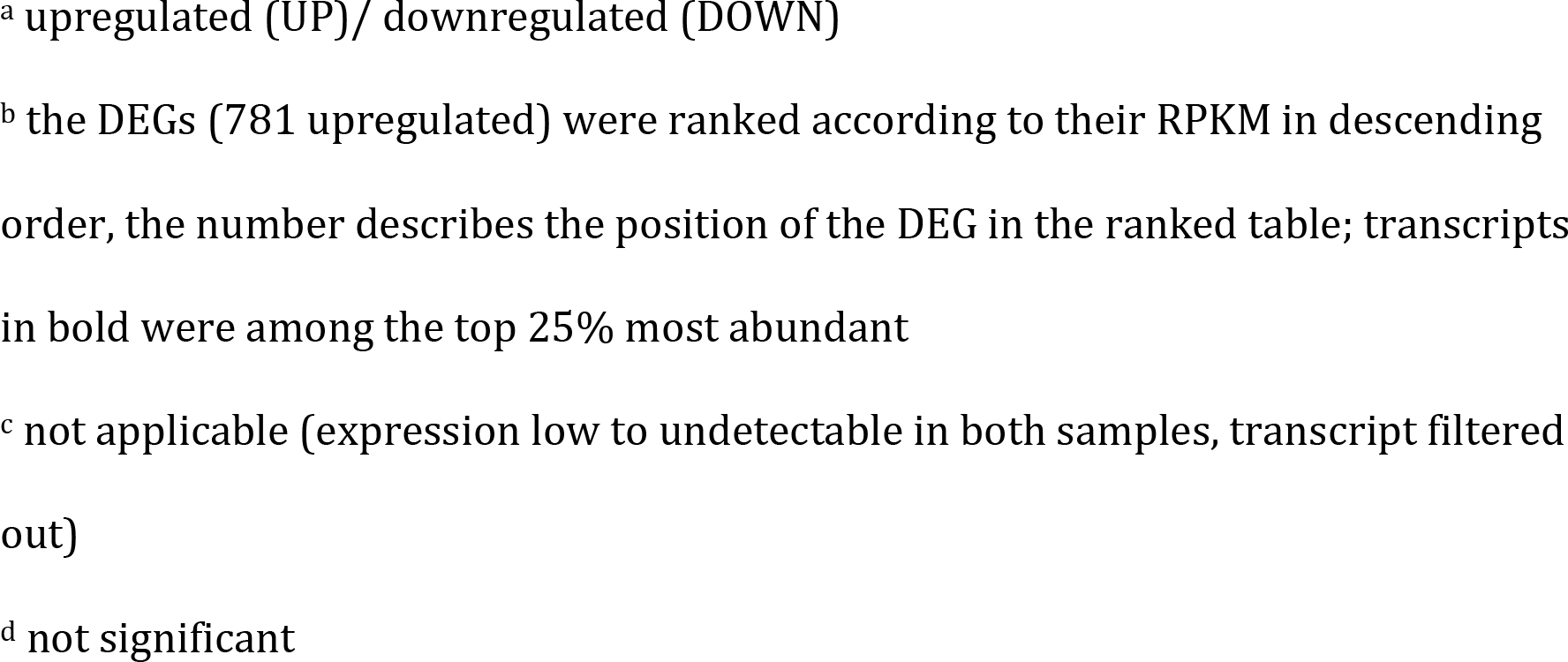

**Figure 7.**
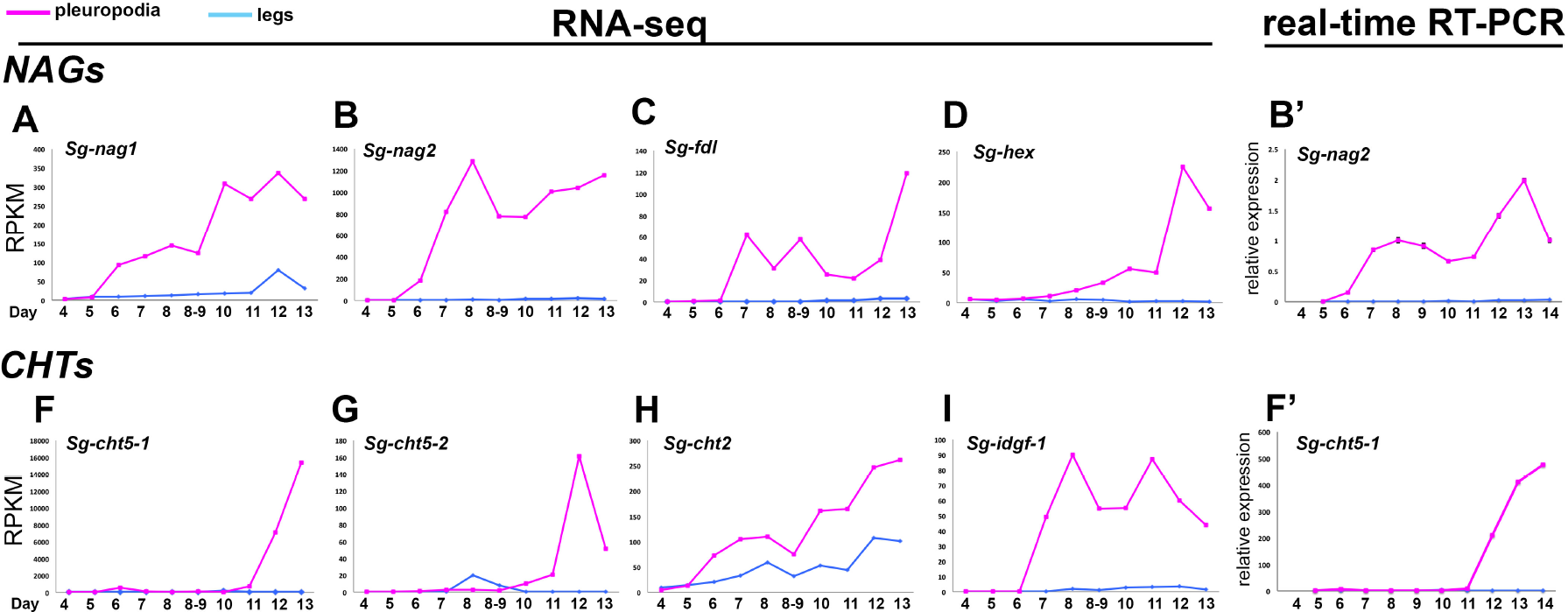
Expression profiles of NAGs and CHTs upregulated in the pleuropodia of *Schistocerca* across development. Top row: NAGs, bottom row: CHTs. (A-D) and (F)-(I): RNA-seq, Expression in single-sample sequencing is shown. (B’) and (F’): real-time RT-PCR. (B’) is the same gene as in (B) and (F’) is the same gene as in (F). Analysis of 3-4 technical replicates is shown. Expression in day 8 was set as 1.

To see if the pleuropodia are the major source of the *Sg-nag2* transcript in the embryo, we looked at its expression in various parts of the body (head, thorax, abdomen with pleuropodia, abdomen from which pleuropodia were removed) using real-time RT-PCR (Figure 8A,B). We performed this analysis in embryos on day 6, when the pleuropodia are still immature, day 8, just at the onset of the secretory activity, day 10 and day 12 during active secretion. During all of the stages the abdomen with pleuropodia had the highest expression (A+ in Figure 8B), although the expression was lower in the youngest sample (day 6) compared to the samples from older embryos (day 8, 10 and 12). This shows that the pleuropodia are the major source of mRNAs for this cuticle-degrading NAG.

**Figure 8.**
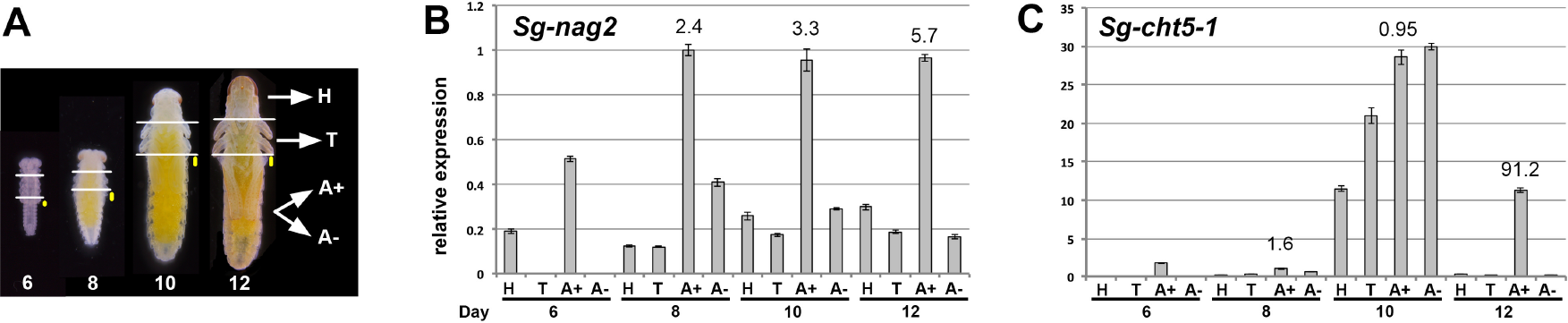
Real-time RT-PCR expression analysis of *Sg-nag2* and *Sg-cht5-1* on cDNA from parts of *Schistocerca* embryos. (A) cDNA was prepared from mRNAs isolated from parts of embryos at the age of 6, 8, 10 and 12 days: H, head; T, thorax; A+, abdomen with pleuropodia; A-, abdomen without pleuropodia. For each age the same number of body parts was used (5-10) and RNA was resuspended in the same volume of water. The size of the pleuropodium is indicated by the yellow dot. (B) and (C): expression of *Sg-nag2* and *Sg-cht5-1*, respectively. Analysis of 3-4 technical replicates is shown. Expression in A+8 (abdomen with pleuropodia at stage when the organs first become differentiated) was set as 1. Numbers above A+ expression is fold change from A- of the same age.

The insect CHTs have been classified into several groups (Zhu et al., 2016; Noh et al., 2018), of which the major role in the digestion of cuticular chitin is played by Chitinase 5 and (perhaps with a secondary importance) by Chitinase 10 (Zhu et al., 2008; Qu et al. 2014) (Table 2; the classification of CHTs into five major groups that we use here is based on Zhu et al., 2008). Some chitinases, for example, are expressed in the gut, trachea and fat body, where they are likely involved in digestion of dietary chitin, turnover of peritrophic matrix and immunity, other chitinases expressed by the epidermis organize assembly of the new cuticle (e.g., Yan et al., 2002; Shi and Paskewitz, 2004; Pesch et al., 2016; Noh et al., 2018).

Our transcriptome contains 16 full or partial transcripts of CHTs representing all of the major CHT groups (Table 2, Figure S5B, S6B). The pleuropodia specifically upregulate both of the genes for Chitinase 5, homologs of *cht5-1* and *cht5-2* from the locust *Locusta migratoria* (Li et al., 2015). One of the transcripts, *Sg-cht5-1*, was among the top 15 most abundant transcripts upregulated in the highly secreting pleuropodia (Table 1). The predicted amino acid sequence contains a conserved catalytic domain and a signal peptide, and thus is likely to be active and secreted, respectively (Figure S5B). The other upregulated CHTs were homologs of *cht2* and *idgf*. By contrast, the *Schistocerca* homolog of *cht10* that also has a role in cuticular chitin hydrolysis and required for larval moulting (Zhu et al., 2008; Pesch et al., 2016) had low expression in both legs and pleuropodia.

We next focused on the transcript of the major chitinase, *Sg-cht5-1*. Unlike the NAGs, both RNA-seq and real-time RT-PCR have shown that the expression of this CHT is low in the early secreting stages, rises only later around day 12 and reaches highest levels when the pleuropodia are already degenerating (day 13 and 14) (Figure 7 F,G,F’). Also real-time RT-PCR on cut embryos has shown that the pleuropodia are a major source of the *Sg-cht5-1* mRNA on day 12 but not before (the high expression in the whole embryo on day 10 could be linked to the second embryonic moult and was also observed with *Sg-cht7*, although not with *Sg-cht10*, Figure S8). These data show that the pleuropodia before hatching express a cuticle-degrading chitinase.

### Pleuropodia upregulate transcripts for some proteases that could digest a cuticle

Our GO enrichment analysis has shown that the secreting pleuropodia are enriched in transcripts for genes associated with proteolysis (Figure 6, Table S11). Transcripts for proteases and their inhibitors are abundant among the top ten percent of the most highly “expressed” upregulated DEGs (Table 1). To see if the upregulated transcripts encode enzymes that are associated with digestion of insect cuticle, we compared our data with the enzymes identified in the complete proteomic analysis of the MF from the lepidopteran *Bombyx mori* (Zhang et al., 2014; Liu et al., 2018). Out of 69 genes that we searched, we found homologs or very similar genes in *Schistocerca* transcriptome for half of them (35). This made in total 75 transcripts, of which 27 were upregulated (seven among the top ten percent most highly expressed) and 15 downregulated (Tables 3, S12). The prominent MF protease Carboxypeptidase A (Sui et al., 2009; Zhang et al., 2014) and the Trypsin-like serine protease known to function in locust moulting (Wei et al., 2007) were not upregulated in the pleuropodia. These data indicate that the pleuropodia upregulate transcripts for proteolytic enzymes associated with the degradation of the cuticle and would be able to contribute to the digestion the SC, although the enzymatic cocktail produced by the pleuropodia may not be identical with the MF.

**Table 3.**
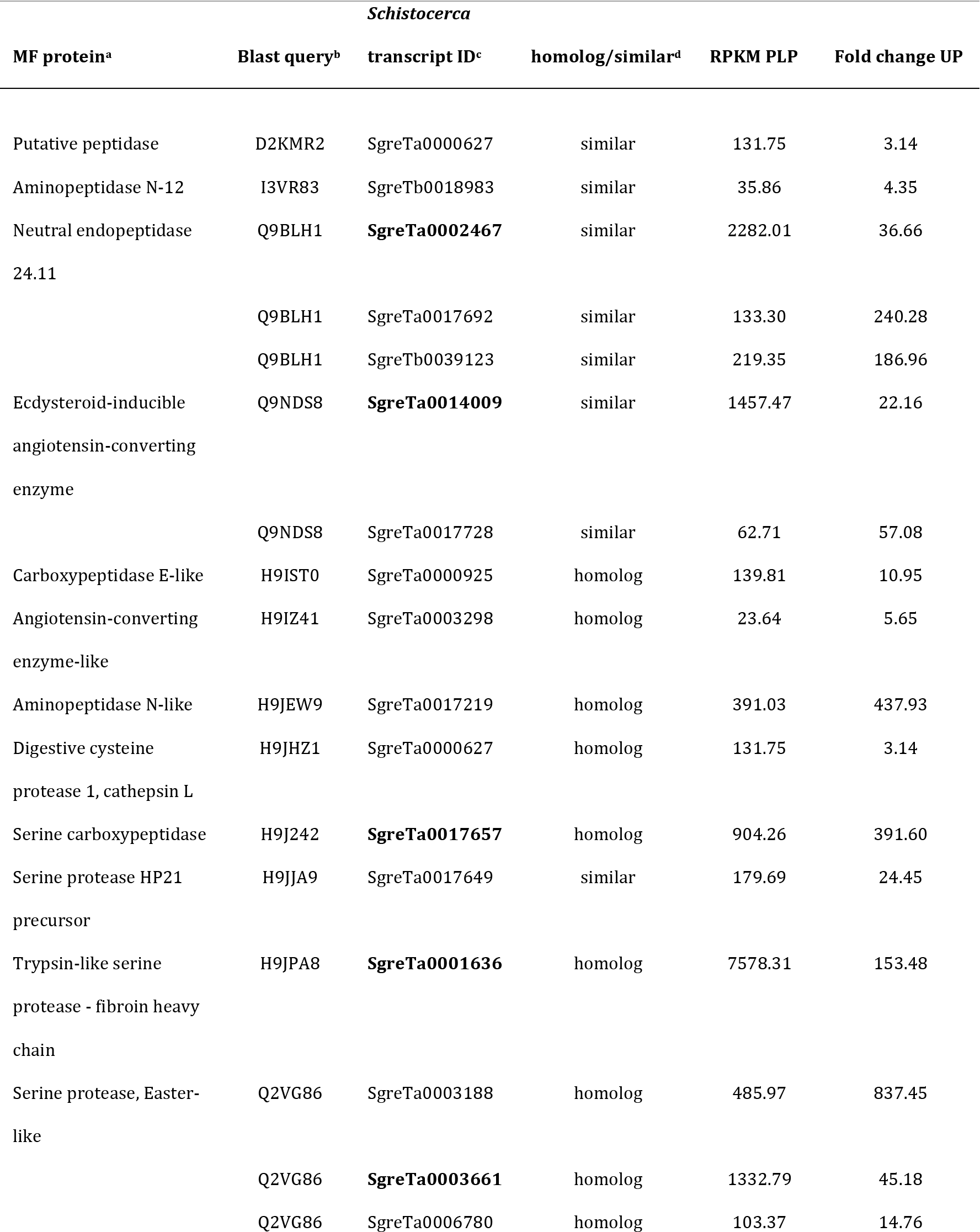
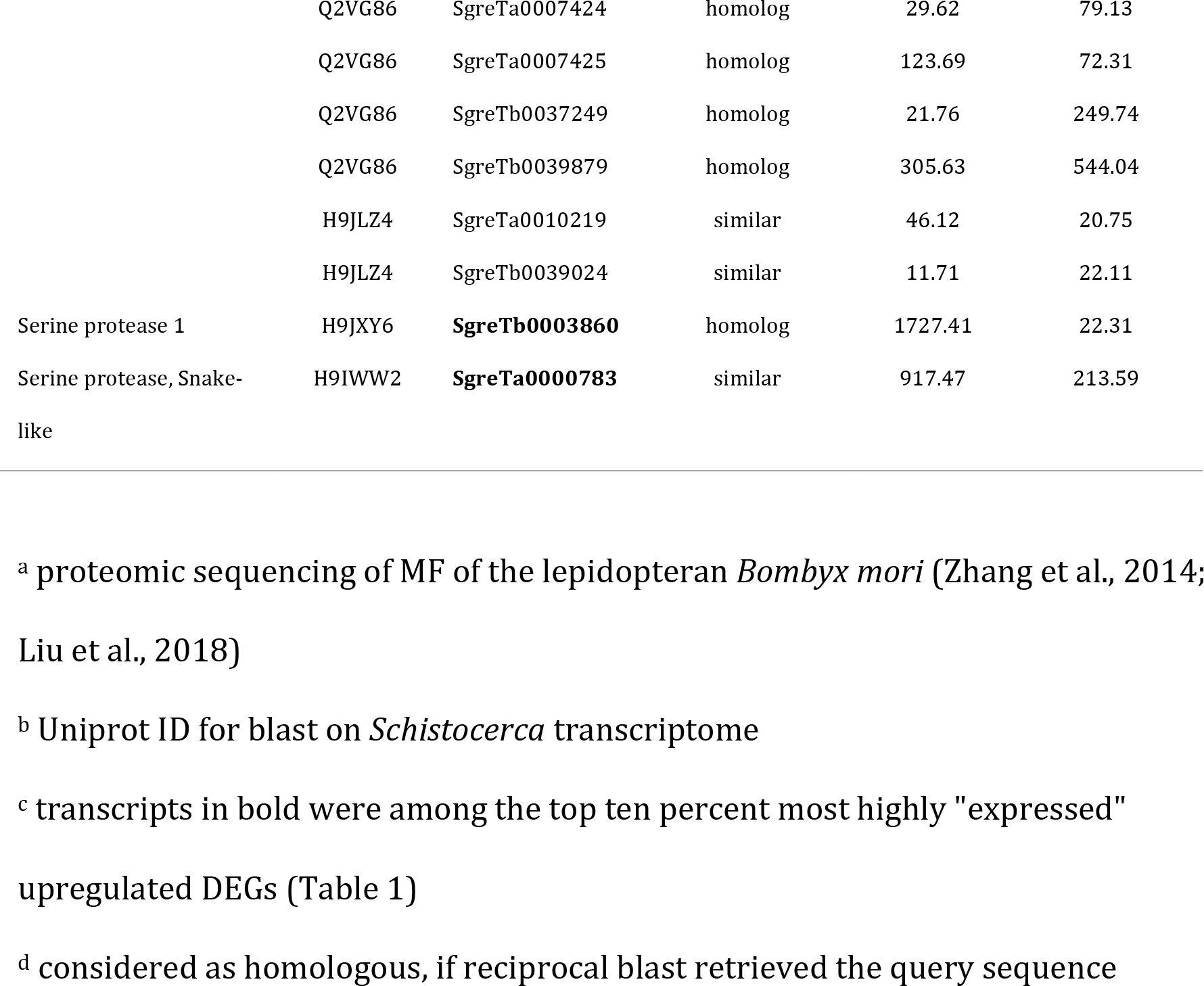
MF proteases that were upregulated in the highly secreting pluropodia of *Schistocerca*.

### Pleuropodia are enriched in transcripts for immunity-related proteins

An observation that was not anticipated was the upregulation of genes for proteins involved in immunity (Lemaitre and Hoffmann, 2007; Buchon et al., 2014) (Figures 6, 9, Table S13). This is especially interesting, because immunity related proteins have been found in the MF (Zhang et al., 2014). This is in agreement with that the cells in the pleuropodia are a type of barrier epithelium (Lemaitre and Hoffmann, 2007; Buchon et al., 2014; Bergman et al., 2017), which enables the contact between the organism and its environment. Barrier epithelia (e.g., the gut, Malpighian tubules or tracheae) constitutively express genes for immune defense.

**Figure 9.**
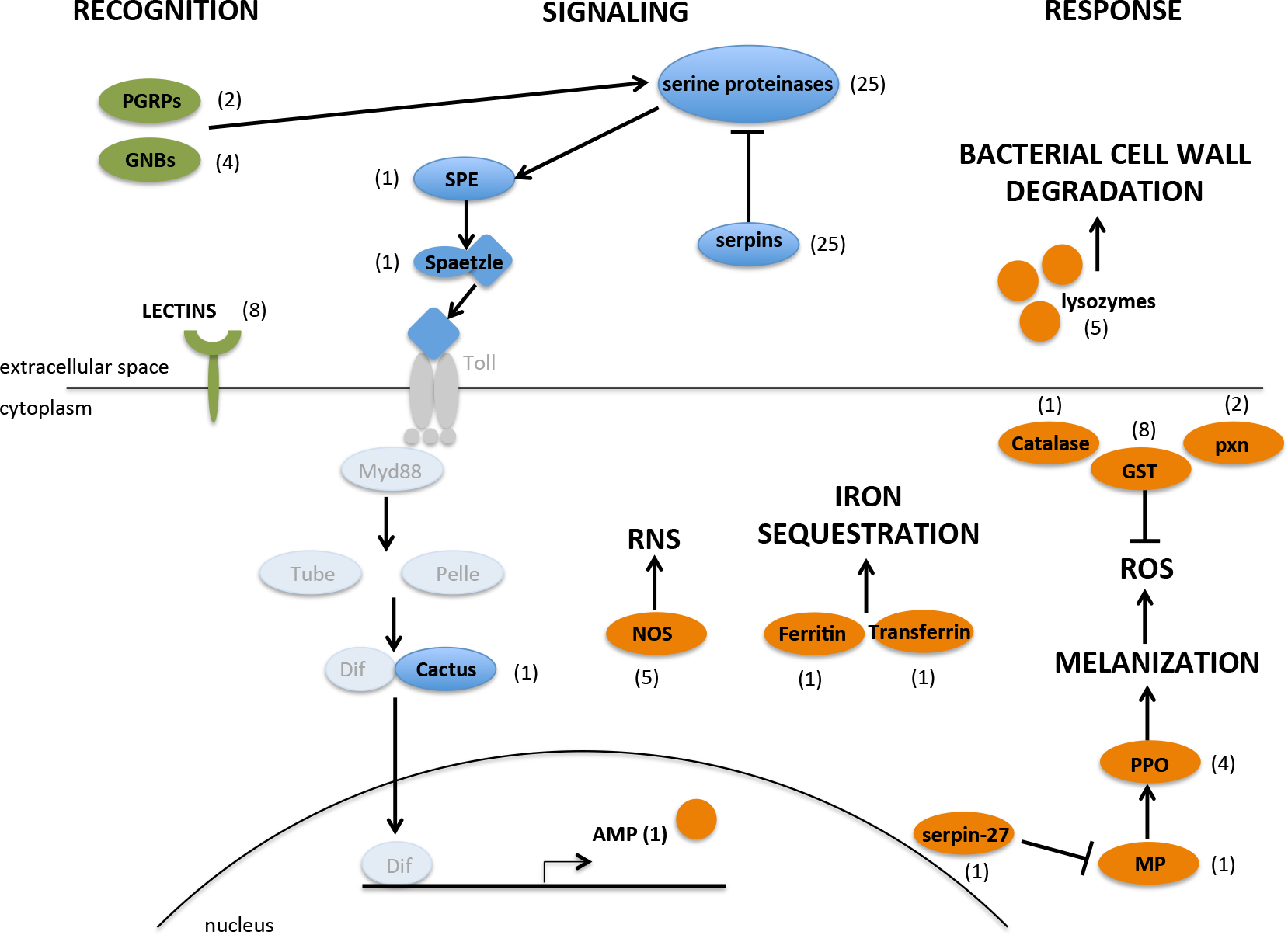
Schematic representation of key immunity-related genes expressed in the highly secreting pleuropodia of *Schistocerca*. Proteins whose transcripts were found in the pleuropodia are in black, number in the brackets is the number of upregulated transcripts. Proteins whose transcripts were not upregulated are in grey. Out of the total 25 serine proteases and 25 serpins, 14 and 15 are known to function in Toll signaling, respectively. AMP, antimicrobial peptide; GNBP, gram-negative bacteria-binding protein; GST, glutathione S-transferase; MP, melanization protease; NOS, nitric oxide synthase; PGRP, peptidoglycan recognition protein; PPO, pro-phenoloxidase; pxn, peroxiredoxin; RNS, reactive nitrogen species; ROS, reactive oxygen species; SPE, Spaetzle-processing enzyme.

**Figure 10.**
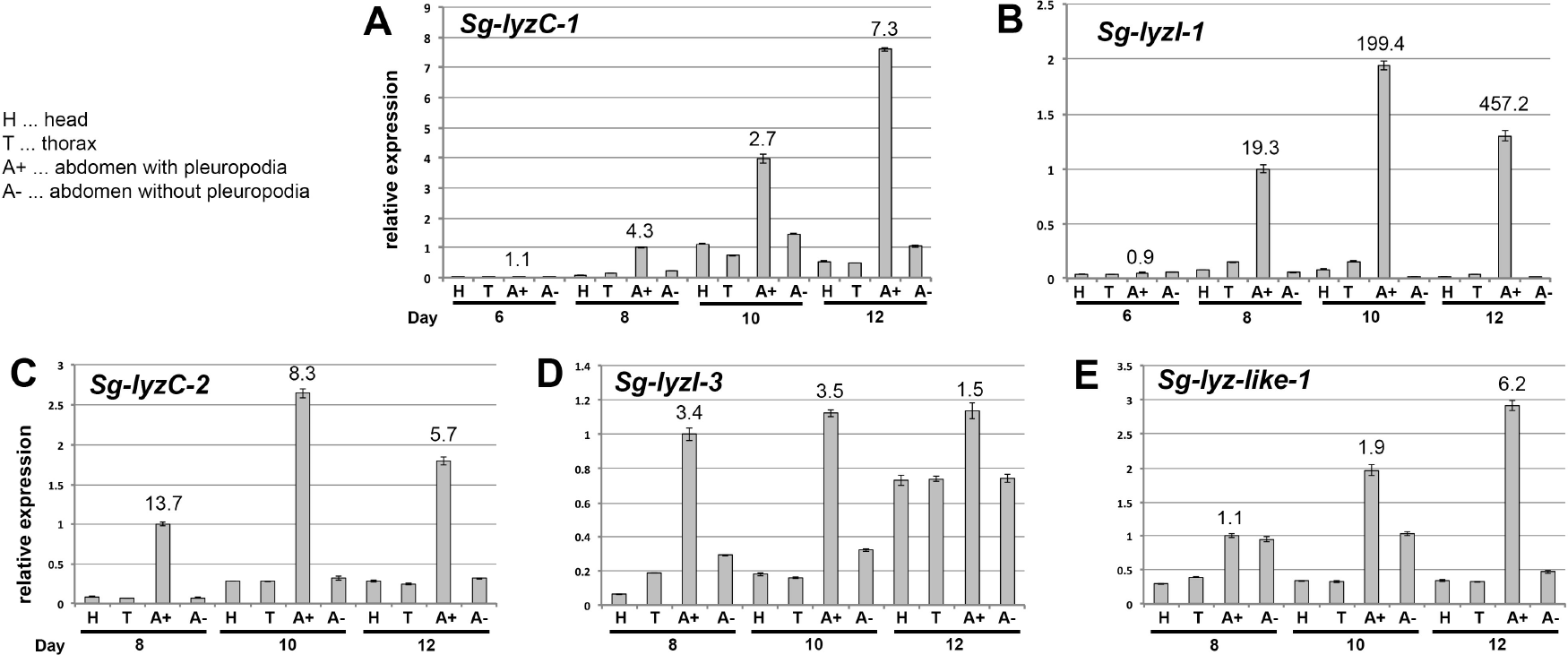
Real-time RT-PCR expression analysis of genes for lysozymes on cDNA from parts of *Schistocerca* embryos. cDNA was prepared from mRNAs isolated from parts of embryos at the age of 6, 8, 10 and 12 days. For each age the same number of body parts was used (5-10) and RNA was resuspended in the same volume of water. Analysis of 3-4 technical replicates is shown. Expression in A+8 (abdomen with pleuropodia at stage when the organs first become differentiated) was set as 1. Numbers above A+ expression is fold change from A- of the same age.

In total we found upregulated 99 transcripts (13 percent of the upregulated genes) for immunity-related proteins. These include proteins at all three levels, the pathogen recognition, signaling and response (Figure 9, Table S13). From the four signaling pathways, Toll was upregulated, but not IMD or JAK/STAT, and from the JNK signaling we found c-Jun. Genes for a range of immune responses were upregulated, including production of reactive nitrogen species (RNS), melanization, genes for lysozymes and one antimicrobial peptide (AMP) similar to Diptericin.

The transcripts for lysozymes were among the most highly expressed (Table 1) and we chose to focus on them. Lysozymes are secreted proteins that kill bacteria by breaking down their cell wall. Our *Schistocerca* transcriptome contains nine genes for lysozymes, seven of which were upregulated (Table 4, Table S14). The second most highly expressed DEG was a transcript for a C-type lysozyme *(Sg-LyzC-1)* that was previously shown to have anti-bacterial properties in *Schistocerca* (Mohamed et al., 2016) (Table 1). We examined expression of five selected genes on cut embryos by real-time RT-PCR (Figure 9). Our data showed that the pleuropodia are the major source of mRNAs for these genes.

**Table 4.** RNA-seq differential gene expression of *Schistocerca* lysozymes in the highly secreting pleuropodia.

| Lysozyme type | Gene | UP/DOWN<sup>a</sup> | Fold change | Expression<sup>b</sup> |
| --- | --- | --- | --- | --- |
| C-type lysozyme | *Sg-LyzC-1* | UP | **336.64** | **2 (0.26%)** |
|  | *Sg-LyzC-2* | UP | **134.71** | **37 (4.74%)** |
| I-type lysozyme | *Sg-LyzI-1* | UP | **550.26** | **7 (0.90%)** |
|  | *Sg-LyzI-2* | ns<sup>c</sup> |  |  |
|  | *Sg-LyzI-3* | UP | **30.62** | **76 (9.73%)** |
|  | *Sg-LyzI-4* | DOWN | -34.41 | 1251 (81.50%) |
|  | *Sg-LyzI-5* | ns |  |  |
| Lysozyme-like | *Sg-Lyz-like-1* | UP | **192.68** | **150 (19.21%)** |
|  | *Sg-Lyz-like-2* | ns |  |  |

<sup>a</sup> upregulated (UP)/ downregulated (DOWN)
<sup>b</sup> the DEGs (781 upregulated) were ranked according to their RPKM in descending order, the number (percentage) describes the position of the DEG in the ranked table; transcripts in bold were among the top 25% most abundant
<sup>c</sup> not significant

### Pleuropodia do not upregulate the pathway for ecdysone biosynthesis

Previous work has suggested that pleuropodia may be embryonic organs producing the moulting hormone ecdysone (Novak and Zambre, 1974). During post-embryonic stages, ecdysone is synthesized in the prothoracic glands and several other tissues by a common set of enzymes (reviewed in Niwa and Niwa 2014; Ou et al., 2016), some which have been characterized in the locusts (Marchal et al., 2011, 2012; Lenaerts et al., 2016; Sugahara et al., 2017). As shown in *Drosophila*, these genes are expressed in diverse cell types in embryos, and when the larval prothoracic glands are formed their expression co-localizes there (Chávez et al., 2000; Warren et al., 2002; Petryk et al., 2003; Niwa et al., 2004; Warren et al., 2004).

Out of the nine genes critical for ecdysone biosynthesis, only one (*dib*) was upregulated in the highly secreting pleuropodia (Table 5, S15). One gene (*spo*) was downregulated. The pleuropodia are not enriched in the whole pathway at any time of development, including around katatrepsis, in which experiments supporting the synthesis of moulting hormone were carried out (Table S9, S16). Under the GO term “hormone biosynthetic process” enriched in the highly secreting pleuropodia (Table S7, S17) we found a gene *Npc2*a that encodes a transporter of sterols including precursors of ecdysone. It is also required for ecdysone biosynthesis, but indirectly and in the cells it functions as a general regulator of sterol homeostasis (Huang et al., 2007). We conclude that our transcriptomic data provide little evidence that the pleuropodia are involved in ecdysone biosynthesis.

**Table 5.** RNA-seq differential gene expression of *Schistocerca* ecdysone biosynthesis enzymes in the highly secreting pleuropodia.

| Gene | UP/DOWN<sup>a</sup> | Fold change | Expression<sup>b</sup> |
| --- | --- | --- | --- |
| *shade (shd), Cyp314A1* | ns<sup>c</sup> |  |  |
| *shadow (sad), Cyp315A1* | ns |  |  |
| *disembodied (dib), Cyp302A1* | UP | 5.71 | 431 (55%) |
| *phantom (phm), Cyp306A1* | ns |  |  |
| *shroud (sro)* | ns |  |  |
| *spook (spo), Cyp307A1* | DOWN | -12.32 | 1368 (89%) |
| *spook-like* | ns |  |  |
| *neverland (nvd)* | ns |  |  |
| *Cyp6t3* | not found |  |  |
| *Cyp6u1* | na<sup>d</sup> |  |  |
| *Cyp303a1* | ns |  |  |

<sup>a</sup> upregulated (UP)/ downregulated (DOWN)
<sup>b</sup> the DEGs (781 upregulated and 1535 downregulated) were ranked according to their RPKM in descending order, the number (percentage) describes the position of the DEG in the ranked table
<sup>c</sup> not significant
<sup>d</sup> not applicable (expression low to undetectable in both samples, transcript filtered out)

## DISCUSSION

### Pleuropodia of *Schistocerca* express genes for the “hatching enzyme”

The first demonstration of the physiological role of the pleuropodia comes from the experiments carried out on a grasshopper *Melanoplus* (closely related to *Schistocerca*), by Eleanor Slifer (Slifer, 1937). When she took embryos before hatching (Figure 2) and separated anterior and posterior halves by ligation, the SC was digested only in the part of the egg with the pleuropodia. Surgical removal of the pleuropodia prevented SC digestion in the whole egg. Slifer’s hypothesis that the pleuropodia secrete the “hatching enzyme” was criticized by Novak and Zambre (1974): if the deposition and digestion of the SC is similar to the cuticle turnover during larval moulting, then the “hatching enzyme” is produced by the serosa. They believed that the pleuropodia reach the peak of their activity in embryos during katatrepsis (45% DT) and participate in digestion of the SC indirectly by secreting ecdysone to stimulate the serosa.

Our ultrastructural observations on staged pleuropodia of *Schistocerca* have shown that the glandular cells only begin to differentiate just at the time of katatrepsis (45% DT) and do not secrete at that time. This would explain why no digestive effect on the SC was detected by Novak and Zambre (Novak and Zambre, 1974) using a homogenate from *Schistocerca* pleuropodia isolated at this stage. The release of granular secretion starts just before dorsal closure (55% DT) and intensifies before hatching. This is in agreement with previous observations on some stages of the pleuropodia in other orthopterans (Louvet, 1975; Viscuso and Sottile, 2008).

Our RNA-seq analysis revealed that the secreting pleuropodia highly express genes encoding enzymes that are capable of digesting a typical chitin-protein insect cuticle. These include genes for proteolytic enzymes similar to those present in the MF and cuticular chitin-degrading NAGs and Chitinase 5. The pleuropodia also express genes for Chitinase 2 and Idgf, which have low effect on cuticular chitin digestion, but were shown to organize proteins and chitin fibres during cuticle deposition (Pesch et al., 2016). These CHTs may organize the fibres in the cuticle secreted by the pleuropodia (Figure 3).

In combination with RT-PCR we showed that, while the expression of the *Sg-nag1* and *Sg-nag2* started to rise in parallel with the differentiation of the glandular cells, the *Sg-cht5-1* and *Sg-cht5-2* transcripts raised shortly before hatching. Chitinase 5 is a critical chitin-degrading chitinase in insects: it is highly abundant in the moulting fluid and its silencing in diverse insects including locusts leads to failure in larval moulting (Zhu et al., 2008; Zhang et al., 2014; Li et al., 2015; Xi et al., 2015; Pesch et al., 2016). Our data indicate that the sudden rise in the expression of *Sg-cht5* in the pleuropodia at the end of embryogenesis and presumably secretion of this CHT into the extraembryonic space is the key component of the “hatching enzyme” effect (Slifer 1937; 1938) in locusts and grasshoppers.

### Pleuropodia in some other insects could secrete the “hatching enzyme” and their function may also vary among species

There is evidence to suggest that the process occurs similarly in some insect. As in orthopterans, the pleuropodia of the rhagophthalmid beetle *Rhagophthalmus ohbai* release secretion after katatrepsis and SC rapidly degrades just shortly before hatching (Kobayashi et al., 2003). In the large water true bugs from the family Belostomatidae, the male carries a batch of eggs on his back. It is believed that the detachment of the eggs just before hatching is also caused by the secretion from the pleuropodia (Tanizawa et al., 2007).

The molecular mechanism of SC degradation may also vary between insects and as previously hypothesized (Novak and Zambre, 1974) the serosa may also contribute to the SC degradation. The serosa of the beetle *Tribolium*, expresses *cht10* and *cht7* (Jacobs et al., 2015), of which the former CHT is important for cuticular chitin digestion. Silencing of *cht10*, but not *cht5* prevented larvae from hatching (Zhu et al., 2008). Transcripts for *cht10* were not upregulated in the pleuropodia of *Schistocerca*. This suggests that the SC is degraded by enzymes produced by both, the serosa and the pleuropodia and that the indispensable roles in cuticle digestion are played by different enzymes in different insects.

In some insects the pleuropodia may not be involved in hatching but have another function. In the viviparous cockroach *Diploptera punctata* (Stay, 1977), the secretion from the pleuropodia is very low and the large pleuropodia of the melolonthid beetle *Rhizotrogus majalis* have not been observed to produce any secretion granules at all (Louvet, 1983). In dragonflies, one of the more basal lineages of insects, the secretion likely has a different function than in orthopterans, because their SC is not digested before hatching (Ando, 1962). The special epithelium in the pleuropodia shares features with transporting epithelia (Louvet, 1973; Stay, 1977) that function in water transport and ion balance (Berridge and Oschman, 1972). Our data do not exclude this function, but it is yet to be tested.

### The pleuropodia of *Schistocerca* are enriched in transcripts for enzymes functioning in immunity

We found that many of the genes expressed in the pleuropodia encode proteins involved in immunity (Lemaitre and Hoffmann, 2007). This indicates that the pleuropodia are also organs of epithelial immunity, similar to other barrier epithelia in postembryonic stages (such as the gut) (Bergman et al., 2017), which are in a constant contact with microorganisms. The pleuropodia differ from such tissues in that they are not directly exposed to the environment, but enclosed in the eggshell, seemingly limiting their contact with microorganisms. Proteins associated with immune defense are also found in the MF (Zhang et al., 2014), where they prevent invasion of pathogens through a “naked” epidermis after the separation of the old cuticle from the epidermis in the process of apolysis. As found in the beetle *Tribolium*, during the early embryonic stages the frontier epithelium providing the egg with an immune defense is the extraembryonic serosa (Jacobs et al., 2014). The serosa starts to degenerate after katatrepsis and disappears at dorsal closure (Panfilio, 2008). The pleuropodia of *Schistocerca* differentiate just before dorsal closure, suggesting that they take over this defense function in late embryogenesis. It will be interesting to clarify in the upcoming research whether apart from their role in hatching the pleuropodia are important organs for fighting against potential pathogens that have gained access to the space between the embryo and the eggshell.

### Conclusions

The pleuropodia of *Schistocerca* have morphological markers of high secretory activity in the second half of embryogenesis after the definitive dorsal closure is finished. Transcriptomic profiling indicate that the conclusions that Eleanor Slifer drew from her experiments over eighty years ago that the pleuropodia secrete cuticle degrading enzymes, were correct. The pleuropodia likely have other functions, such as in immunity. The pleuropodia are specialized embryonic organs and apparently an important though neglected part of insect physiology.

## MATERIALS AND METHODS

### Insects

*Schistocerca gregaria* (gregarious phase) were obtained from a long-term, partly inbred colony at the Department of Zoology, University of Cambridge. Eggs were collected into aluminium pots filled with damp sand. The pots were picked up after two (most samples) or four hours and incubated at 30°C.

### Description of embryonic stages

Embryos and appendages were dissected in phosphate buffer saline (PBS). Whole eggs were bleached in 50% household bleach to dissolve the chorion. All were photographed in water or PBS using the Leica M125 stereomicroscope equipped with DFC495 camera and associated software. Photos were processed using Adobe Photoshop CC 2017.1.1. Photos of eggs and embryos that illustrate the stage (Figure 2A and S1) had the background cleaned using the software (removal of the tools that hold the photographed objects in place).

### Immunohistochemistry on paraffin sections

Embryos were dissected in PBS and pieces including posterior thorax and anterior abdomen (older embryos) or mid thorax plus whole abdomen (young embryos) were fixed in PEMFA (4% formaldehyde in PEM buffer: 100 mM PIPES, 2.0 mM EGTA, 1.0 mM MgSO4) at room temperature (RT) for 15-30 minutes, then washed in PBT (PBS with 0.1% Triton-X 100) and stored in ethanol at −20°C. Samples were cleared in 3×10 minutes in Histosol (National Diagnostics) at RT, infiltrated with paraffin at 60°C for 2-3 days, embedded in moulds and hardened at RT. Sections 6-8 μm thick were prepared on a Leica RM2125RTF microtome. The slides with sections were washed with Histosol, ethanol, then stepwise re-hydrated to PBT. Incubations were carried out in a humidified chamber. Slides were blocked with 10% sheep serum (Sigma-Aldrich) in PBT for 30 minutes at RT, incubated with Phospho-Histone H3 antibody (Invitrogen) diluted with PBT 1:130 at 4°C overnight, washed and incubated with Alexa Fluor 568 anti-rabbit secondary antibody (Invitrogen) diluted 1:300 at RT for 2 hours, washed and incubated with DAPI (Invitrogen) diluted 1:1000. Sections were imaged with a Leica TCS SP5 confocal microscope and photos processed using Fiji (https://fiji.sc).

### Electron microscopy

For TEM embryos were removed from the chorion in PBS and pieces of posterior thorax to anterior abdomen were fixed in 2.5-3.0% glutaraldehyde in 0.1 M phosphate buffer pH7.2 for a few hours at room temperature and then at 4°C for several days. Each pleuropodium and leg were then separated and embedded into 2% agar. Small cubes of agar with the tissue were incubated in osmium ferrocyanide solution (3% potassium ferricyanide in cacodylate buffer with 4 mM calcium chloride) for 1-2 days at 4°C, then in thiocarbohydrazide solution (0.1 mg thiocarbohydrazide from Sigma-Aldrich, and 10 ml deionized water dissolved at 60°C) and protected from light for 20-30 minutes at RT, then in 2% aqueous osmium tetroxide 30-45 minutes at RT and in 1% uranyl acetate (maleate buffered to pH 5.5) at 4°C overnight. Washing between each step was done with deionized water. Samples were dehydrated in ethanol, washed with dry acetone, dry acetonitrile, infiltrated with Quetol 651 resin (Agar Scientific) for 4-6 days and hardened in moulds at 60°C for 2-3 days. Semithin sections were stained with toluidine blue. Ultrathin sections were examined in the Tecnai G280 microscope. For SEM whole embryos were dissected out of the chorion in PBS, fixed in 3% glutaraldehyde in phosphate buffer similarly as above. They were post-fixed with osmium tetroxide, dehydrated through the ethanol series, critical point dried, gold coated, and observed in a FEI/Philips XL30 FEGSEM microscope. Photos from TEM and SEM were processed using Adobe Photoshop CC 2017.1.1.

### Preparation of the reference transcriptome

Whole embryo transcriptome: Eggs from each 1-day egg collection incubated for the desired time were briefly treated with 50% bleach, washed in distilled water and frozen in liquid nitrogen. Total RNA was isolated with TRIzol reagent (Invitrogen), treated with TURBO DNase (Invitrogen) and purified on a column supplied with the RNAeasy Kit (Quiagen). The purified RNA from each day (14 samples) was pooled into 4 samples: day 1-4, 5-7, 8-10 and 11-14. Ten μg of RNA from each of the 4 samples was sent to BGI (Hong Kong). The total RNA was enriched in mRNA by using the oligo(dT) magnetic beads and cDNA library was prepared. 100 bp paired-end (PE) reads were sequenced on Illumina HiSeq 2000; numbers of the reads obtained are in Table S2. Non-clean reads were filtered using filter_fq software (removes reads with adaptors, reads with unknown nucleotides larger than 5% and low quality reads). Transcripts from all samples were assembled separately using the Trinity software (release 20130225) (Grabherr et al., 2011) with parameters: --seqType fq --min_contig_length 100; --min_glue 4 --group_pairs_distance 250; -- path_reinforcement_distance 95 --min_kmer_cov 4. Transcriptes from the 4 assemblies were then merged together to form a single set of non-redundant transcripts using TGICL software (v2.1) (Pertea, 2003) with parameters: -l 40 -c 10 -v 20.

Legs and pleuropodia transcriptome (age about 8.5-8.75 days): The appendages were dissected in cold RNase-free PBS (treated with diethyl pyrocarbonate) and total RNA was isolated and cleaned as described above. Ten μg of RNA from each leg sample and pleuropodium sample were transported to the Eastern Sequence and Informatics Hub (EASIH), Cambridge (UK). cDNA libraries were prepared including mRNA enrichment. 75 bp PE reads were sequenced on Illumina GAIIX; numbers of the reads obtained are in Table S2. The reads were trimmed to the longest contiguous read segment for which the quality score at each base is greater than a Phred quality score of Q = 13 (or 0.05 probability of error) using the program DynamicTrim (v. 1.7) from the package SolexQA (Cox et al., 2010; http://solexaqa.sourceforge.net/). The trimmed reads were then filtered to remove sequence adapter using the program cutadapt (v. 0.9; http://code.google.com/p/cutadapt/). Sequences shorter than 40 bp were discarded. Trimmed reads were used to de novo assemble the transcriptome using Velvet (v. 1.1.07; Zerbino et al., 2008; http://www.ebi.ac.uk/~zerbino/velvet/) (commands: -shortPaired –fastq; -short2 –fastq; -read_trkg yes) and Oases (v.0.2.01; Schulz et al., 2012; http://www.ebi.ac.uk/~zerbino/oases/) (commands: - ins_length 350). Velvet is primarily used for de-novo genome assembly; here, the contigs that were output by Velvet were used by the complementary software package Oases to build likely transcripts from the RNA-seq dataset. K-mer sizes of 21, 25 and 31 were attempted for the two separate samples as well as the combined samples and optimal K-mer sizes of 21 were found for both samples.

Transcripts for the reference transcriptome were selected from the embryonic and legs and pleuropodia transcriptomes. The transcripts were first merged with evigene (version 2013.03.11) using default parameters. Because this selection of transcripts eliminated some genes (gene represented by zero transcripts, although the transcripts were present in the original transcriptomes), we repeated the step with less strict parameters (cd-hit-est - version 4.6, with -c 0.80 -n 5). This second selection contained several genes represented by more transcripts, therefore we aligned selection 1 and 2 to each other to identify, which genes in selection 1 were missing. Selection 1 was then completed with the help of selection 2 by adding the missing transcripts. The quality and completeness of the resulting transcriptome was assessed and edited in the following steps. First, we removed several redundant transcripts manually: these were found by blasting diverse insect sequences (queries) against the *Schistocerca* transcriptome using the local ViroBLAST interface (Deng et al., 2007). Some transcripts were edited manually, such as when we found that two transcripts were combined into one, resulting in an alignment against two protein sequences (*Schistocerca* transcript blasted against NCBI GenBank database) we split the respective transcripts. Second, we blasted the whole transcriptome against itself and removed redundant sequences, if the alignment was spanning at least 300 bp with a sequence identity of at least 98% (Blast+ suite, version 2.6.0). The longer transcript was kept in all cases. Transcripts shorter than 200 bp were discarded. All these steps were carried out in R (R Development Core Team, 2008; http://www.R-project.org) and sequences were handled using the Biostrings package (Pagès et al., 2017).

### Sequence analysis

Basic transcript analysis was done by CLC Sequence Viewer7 (QIAGEN). Signal peptide and transmembrane regions were predicted by Phobius (Käll et al., 2007; http://phobius.binf.ku.dk/index.html). Conserved domains were identified using SMART (http://smart.embl-heidelberg.de/). To annotate the newly assembled transcriptome, the freely available annotation pipeline Trinotate (version 3.1.1) was used (Haas et al., 2013). The longest candidate ORF of each sequence was identified with the help of the inbuilt TransDecoder (Haas et al., 2013; https://github.com/TransDecoder/TransDecoder/wiki) software.

A blast was run against Uniprot sequences specific for *Schistocerca gregaria, Locusta migratoria, Apis melifera, Tribolium castaneum, Bombyx mori* and *Drosophila melanogaster* (blastx with default parameter and -max_target_seqs 1).

### RNA-seq expression analysis

Pleuropodia and hind legs from embryos at the same age (day 4, 5, 6, 7, 8, 10, 11, 12 and 13) were dissected in cold RNase-free PBS and total RNA was isolated as described for samples for the reference transcriptome, but cleaned with RNA Clean & Concentrator (Zymo Research). One μg of RNA from each sample was sent to BGI (Hong Kong). The mRNA enrichment and cDNAs preparation was as described above. 50 bp single-end (SE) reads were sequenced on Illumina HiSeq 2000. Over 45 million reads were sequenced from each sample (Table S2).

A pair of samples from mixed embryos 8-9 days that was used for the preparation of the reference transcriptome (described above) was also included in the expression analysis, but prior to mapping, the 75 bp PE reads were trimmed to 50 bp, using Trimmomatic in the paired-end mode (version 0.36) using the CROP function (CROP:50). A single pleuropodium or leg sample was sequenced from each stage.

The quality of the sequenced reads was assessed using the FastQC software. All samples consistently showed a Per base sequence quality of > 30. Reads were mapped to the reference transcriptome with Bowtie2 (version 2.2.5) using default parameter and the –local alignment mode (Langmead et al., 2009). The trimmed pairs of reads were concatenated for each stage and treated as single reads. A PCA plot was generated to assess if differences in sequencing type and processing (SE samples and PE samples day 8-9) had an effect, which was not the case. This plot was prepared by using the plotPCA() function in the DESeq2 R package (Love et al., 2014); the count matrix was transformed with the rlog() function. The R package HTSFilter (Rau et al., 2013) was used with default parameters to filter constantly low expressed genes and 12988 transcripts were left.

The differential expression analysis was performed with the NOISeq R package (2.22.1; Tarazona et al., 2011). Reads were first normalized using the RPKM method (Mortazavi et al., 2008). We used NOISeq-sim to find the differentially expressed genes between legs and pleuropodium for each stage with the following parameters: k = NULL, norm =“n”, pnr =0.2, nss =5, v = 0.02, lc=1, replicates =“no”, following the recommendations by the authors for simulation of “technical replicates” prior to differential expression analysis without replicates. Additionally differentially expressed genes between highly secreting pleuropodia and legs at the same stage were assessed (treating samples from day 10, 11 and 12 as replicates) using the NOISeq-real algorithm with the following parameters: k=0.5, norm=“n”, factor=“type”, nss=0, lc=1, replicates = “technical”. To define significantly differentially expressed genes, the probability (“prob”) threshold was set at 0.7 for single stage comparisons and 0.8 for the triplicated comparison, RPKM > 10 and fold change > 2 for both single stage and triplicated comparisons (based on the expression of the genes whose expression dynamics in the pleuropodia were already known, Table S4).

### GO enrichment

The transcriptome was blasted against the whole UniProt/Swiss-Prot database to assess the corresponding GO terms. Only blast hits with an e-value <= 1e-5 were considered for the subsequent GO annotation. GO enrichment of differentially expressed genes was performed using the R package GOSeq (version 1:30.0, Young et al., 2010) implemented in the Trinotate pipeline (see above). Enriched GO terms were summarized and visualized with REVIGO (Supek et al., 2011). Dot plots were prepared from DEGs selected at thresholds: RPKM > 50, fold change > 3.

### Real-time RT-PCR

Tissues were dissected, total RNA was isolated and DNase treated the same way as for sequencing and cleaned with RNA Clean & Concentrator (Zymo Research). cDNA was synthesized with oligo-dT primer (Invitrogen) and 0.5 μg (legs, pleuropodia) or 1 μg (pieces of embryos) of the RNA using ThermoScript RT-PCR System (Invitrogen) at 55°C. The cDNA was diluted to concentration 40 ng/μl and 5 μl was used in a reaction containing 10 μl of SYBR Green PCR Master Mix (Applied Biosystems) and 5 μl of a 1:1 mix of forward and reverse primers (each 20 nM in this mix). Reactions were run in the LightCycler480 (Roche) and analyzed using the associated software (release 1.5.0 SP1) according to the comparative Ct method and normalized to the *eEF1α* gene. Primers (Table S18) were designed with Primer3PLUS program (Untergasser et al., 2007). To check for the presence of a single PCR product, the melting curve was examined after each run and for each pair of primers at least 2 finished runs were visualized on a 2% agarose gel.

The program was: denaturation: 95°C for 10 minutes (1 cycle), amplification: 95°C for 10 seconds, 60°C for 15 seconds, 72°C for 12 seconds (40 cycles) melting: 95°C for 5 seconds, 60°C for 1 minute, 95°C.

## LIST OF ABBREVIATIONS

A1: first abdominal segment
CHT: chitinase
DEG: differentially expressed gene
DT: developmental time
EC1, EC2, EC3: the first, the second, the third embryonic cuticle, respectively
GO: gene ontology
LEG: hind leg(s)
MF: moulting fluid
NAG: β-N-acetyl-hexosaminidase
PCA: principal component analysis
PLP: pleuropodium (pleuropodia)
RPKM: reads per kilobase of transcript per million reads mapped
SC: serosal cuticle
SEM: scanning electron microscopy
T3: third thoracic segment
TEM: transmission electron microscopy

## AUTHOR’S CONTRIBUTIONS

BK initiated the study, designed research, carried out all experimental work, supervised the bioinformatics analysis, interpreted the data and wrote the paper; EB performed majority of the bioinformatics analysis and edited the draft; AC carried out the initial steps in the selections of transcripts for the reference transcriptome and did a preliminary expression analysis. All authors read and approved the manuscript.

## ACKNOWLEDGEMENTS

Majority of the work was carried out in the lab of Michael Akam (University of Cambridge) and the data analysis was finished in the lab of Gregor Bucher (University of Göttingen); BK thanks to both for hosting and financial support. Electron microscopy was done at the Cambridge Advanced Imaging Centre (University of Cambridge). Immunolabeling was done in the lab and with help of Andrew Gillis. Stereomicroscopic pictures were taken in the lab of Paul Brakefield. We also thank for help and advice to Ken Siggens, Jenny Barna, Jeremy Skepper and lab, Steven Van Belleghem, Barry Denholm, Jan Sobotnik, and Gareth Griffiths, for scripts to Erik Clark and Simon Martin. We thank to Michael Akam, Siegfried Roth, Stuart Reynolds, Nico Posnien and Maurijn van der Zee for comments on the manuscript.

## REFERENCES

Ando H. 1962. The comparative embryology of Odonata with special reference to a relic dragonfly Epiophlebia superstes Selys. Sugadaira Biological Laboratory of Tokyo Kyoiku University.

Ando H, Haga K. 1974. Studies on the Pleuropodia of Embioptera, Thysanoptera, and Mecoptera. Tokyo U. Educ. Sugadaira Biol. Lab. Bull 6:1–8.

Angelini DR, Liu PZ, Hughes CL, Kaufman TC. 2005. Hox gene function and interaction in the milkweed bug Oncopeltus fasciatus (Hemiptera). Dev Biol 287:440–455.

Bedford GO. 1978. The development of the egg of Didymuria violescens (Phasmatodea: Phasmatidae: Podacanthinae) – embryology and determination of the stage at which first diapause occurs. Aust J Zool 18:155–169.

Bennett RL, Brown SJ, Denell RE. 1999. Molecular and genetic analysis of the Tribolium Ultrabithorax ortholog, Ultrathorax. Dev Genes Evol 209:608–619.

Bergman P, Seyedoleslami Esfahani S, Engström Y. 2017. Drosophila as a Model for Human Diseases-Focus on Innate Immunity in Barrier Epithelia. Curr Top Dev Biol 121:29–81.

Bernays EA. 1971. The vermiform larva of *Schistocerca gregaria* (Forskål): form and activity (Insecta, Orthoptera). Z Morph Tiere 70:183–200.

Berridge MJ, Oschman JL. 1972. Transporting epithelia. Academic Press, New York.

Buchon N, Silverman N, Cherry S. 2014. Immunity in Drosophila melanogaster – from microbial recognition to whole-organism physiology. Nat Rev Immunol 14:796–810.

Bullière F. 1970. L’évolution des pleuropodes au cours du développement embryonnaire de la Blabera craniifer (Insecte Dictyoptère). Arch Anat Microsc 59:201–220.

Chávez VM, Marqués G, Delbecque JP, Kobayashi K, Hollingsworth M, Burr J, Natzle JE, O’Connor MB. 2000. The Drosophila disembodied gene controls late embryonic morphogenesis and codes for a cytochrome P450 enzyme that regulates embryonic ecdysone levels. Development 127:4115–4126.

Chintapalli VR, Wang J, Herzyk P, Davies SA, Dow JA. 2013. Data-mining the FlyAtlas online resource to identify core functional motifs across transporting epithelia. BMC Genomics 14:518.

Cox MP, Peterson DA, Biggs PJ. 2010. SolexaQA: At-a-glance quality assessment of Illumina second-generation sequencing data. BMC Bioinformatics 11:485.

Deng W, Nickle DC, Learn GH, Maust B, Mullins JI. 2007. ViroBLAST: a stand-alone BLAST web server for flexible queries of multiple databases and user’s datasets. Bioinformatics 23:2334–2336.

Fraulob M, Beutel RG, Machida R, Pohl H. 2015. The embryonic development of Stylops ovinae (Strepsiptera, Stylopidae) with emphasis on external morphology. Arthropod Struct Dev 44:42–68.

Fukamizo T, Kramer KJ. 1985. Mechanism of chitin oligosaccharide hydrolysis by the binary enzyme chitinase system in insect moulting fluid. Insect Biochem 15:1–7.

Goltsev Y, Rezende GL, Vranizan K, Lanzaro G, Valle D, Levine M. 2009. Developmental and evolutionary basis for drought tolerance of the Anopheles gambiae embryo. Dev Biol 330:462–470.

Graber V. 1889. Ueber den Bau und die phylogenetische Bedeutung der embryonalen Bauchänhange der Insekten. Biol Zent Bl 9:355–363.

Grabherr MG, Haas BJ, Yassour M, Levin JZ, Thompson DA, Amit I, Adiconis X, Fan L, Raychowdhury R, Zeng Q, Chen Z, Mauceli E, Hacohen N, Gnirke A, Rhind N, di Palma F, Birren BW, Nusbaum C, Lindblad-Toh K, Friedman N, Regev A. 2011. Full-length transcriptome assembly from RNA-Seq data without a reference genome. Nat Biotechnol 29:644–652.

Grellet P. 1971. Variations du volume et teneur en AND des noyaux de Scapsipedus marginatus Afz. et Br. (Orthoptère, Gryllidae) au cours de l’embryogenèse. Wilhelm Roux Arch Entwickl Mech Mech Org, 167:243–265.

Haas BJ, Papanicolaou A, Yassour M, Grabherr M, Blood PD, Bowden J, Couger MB, Eccles D, Li B, Lieber M, MacManes MD, Ott M, Orvis J, Pochet N, Strozzi F, Weeks N, Westerman R, William T, Dewey CN, Henschel R, LeDuc RD, Friedman N, Regev A. 2013. De novo transcript sequence reconstruction from RNA-seq using the Trinity platform for reference generation and analysis. Nat Protoc 8:1494–1512.

Hagan HR. 1931. The embryogeny of the polyctenid, Hesperoctenes fumarius Westwood, with the reference to viviparity in insects. J Morphol Physiol 51:3–115.

Heming BS. 1993. Origin and fate of pleuropodia in embryos of Neoheegeria verbasci (Osborn) (Thysanoptera: Phlaeothripidae). In: Bhatti JS, editor. Advances in Thysanopterology. New Delhi: Sciantia Publishing. Journal of Pure and Applied Zoology 4. p 205–223.

Hogenkamp DG, Arakane Y, Kramer KJ, Muthukrishnan S, Beeman RW. 2008. Characterisation and expression of β-N-acetylhexosaminidase gene family of Tribolium castaneum. Insect Biochem Mol Biol 38:478–489.

Huang X, Warren JT, Buchanan J, Gilbert LI, Scott MP. 2007. Drosophila Niemann-Pick type C-2 genes control sterol homeostasis and steroid biosynthesis: a model of human neurodegenerative disease. Development 134:3733–3742.

Hughes CL, Kaufman TC. 2002. Hox genes and the evolution of the arthropod body plan. Evol Dev 4:459–99.

Hussey PB. 1926. Studies on the Pleuropodia of Belostoma Flumineum Say and Ranatra Fusca Palisot de Beauvois, with a discussion of these organs in other insects. Entomol Am 7:1–82.

Jacobs CG, Spaink HP, van der Zee M. 2014. The extraembryonic serosa is a frontier epithelium providing the insect egg with a full-range innate immune response. eLife 3:e04111.

Jacobs CGC, Braak N, Lamers GEM, van der Zee M. 2015. Elucidation of the serosal cuticle machinery in the beetle Tribolium by RNA sequencing and functional analysis of Knickkopf1, Retroactive and Laccase2. Insect Biochem Mol Biol 60:7–12.

Jones B. 1956. Endocrine activity during insect embryogenesis. Control of events in development following the embryonic moult (Locusta migratoria and Locustana pardalina, Orthoptera). J Exp Biol 33:685–696.

Käll L, Krogh A and Sonnhammer ELL. 2007. Advantages of combined transmembrane topology and signal peptide prediction-the Phobius web server. Nucleic Acids Res 35:W429–432.

Kamiya A, Ando H. 1985. External morphogenesis of the embryo of Ascalaphus ramburi (Neuroptera, Ascalaphidae). In: Recent Advances in Insect Embryology in Japan. H Ando, K Miya, editors. (ISEBU Co. Ltd.), p 203–213.

Kjer KM, Simon C, Yavorskaya M, Beutel RG. 2016. Progress, pitfalls and parallel universes: a history of insect phylogenetics. J R Soc Interface 13:20160363.

Kobayashi Y, Ando H. 1990. Early embryonic development and external features of developing embryos of the caddisfly, Nemotaulius admorsus (Trichoptera: Limnephilidae). J Morphol 203:69–85.

Kobayashi Y, Suzuki H, Ohba N. 2003. Development of the pleuropodia in the embryo of the glowworm Rhagophthalmus ohbai (Rhagophthalmidae, Coleoptera, Insecta), with comments on their probable function. Proc Arthropod Embryol Soc Jpn 38:19–26.

Konopova B, Zrzavy J. 2005. Ultrastructure, development, and homology of insect embryonic cuticles. J Morphol 264:339–362.

Lambiase S, Grigolo A, Morbini P. 2003. Ontogenesis of pleuropodia in defferent species of Blattaria (Insecta): a comparative study. Ital J Zool 70:205–212.

Langmead B, Trapnell C, Pop M, Salzberg SL. 2009. Ultrafast and memory-efficient alignment of short DNA sequences to the human genome. Genome Biol 10:R25.

Larink O. 1983. Embryonic and postembryonic development of Machilidae and Lepismatidae (Insecta: Archaeognatha). Entomol Gen 8:119–133.

Lemaitre B, Hoffmann J. 2007. The host defense of Drosophila melanogaster. Annu Rev Immunol 25:697–743.

Lenaerts C, Van Wielendaele P, Peeters P, Vanden Broeck J, Marchal E. 2016. Ecdysteroid signalling components in metamorphosis and development of the desert locust, Schistocerca gregaria. Insect Biochem Mol Biol 75:10–23.

Lewis DL, DeCamillis M, Bennett RL. 2000. Distinct roles of the homeotic genes Ubx and abd-A in beetle embryonic abdominal appendage development. Proc Natl Acad Sci USA 97:4504–4509.

Li D, Zhang J, Wang Y, Liu X, Ma E, Sun Y, Li S, Zhu KY, Zhang J. 2015. Two chitinase 5 genes from Locusta migratoria: molecular characteristics and functional differentiation. Insect Biochem Mol Biol 58:46–54.

Liu HW, Wang LL, Tang X, Dong ZM, Guo PC, Zhao DC, Xia QY, Zhao P. 2018. Proteomic analysis of Bombyx mori molting fluid: Insights into the molting process. J Proteomics 173:115–125.

Locke M, Krishnan N. 1973. The formation of the ecdysial droplets and the ecdysial membrane in an insect. Tissue Cell 5:441–450.

Louvet JP. 1973. L’ultrastructure du pleuropode et son ontogenèse, chez l’embryon du phasme Carausius morosus Br. I. – Étude du pleuropode de l’embryon agé. Ann Sci Nat Zool 12:525–594.

Louvet JP. 1975. Premières observations sur l’ultrastructure du pleuropode chez le Criquet migrateur. C R Acad Sci Paris D 280: 1301–1304.

Louvet JP. 1983. Ultrastrucutre du pleuropode chez l’embryon du hanneton Rhizotrogus majalis Razoum (Coleoptera: Melolonthidae). Int J Insect Morphol Embryol 12:97–117.

Love MI, Huber W, Anders S. 2014. Moderated estimation of fold change and dispersion for RNA-seq data with DESeq2. Genome Biol 15:550.

Machida R. 1981. External features of embryonic development of a jumping bristletail, Pedetontus unimaculatus Machida (Insecta, Thysanura, Machilidae). J Morphol 168:339–355.

Machida R, Tojo K, Tsutsumi T, Uchifune T, Klass K-D, Picker MD, Pretorius L. 2004. Embryonic development of heel-walkers: reference to some prerevolutionary stages (Insecta: Mantophasmatodea). Proc Arthropod Embryol Soc Jpn 39:31–39.

Marchal E, Badisco L, Verlinden H, Vandersmissen T, Van Soest S, Van Wielendaele P, Vanden Broeck J. 2011. Role of the Halloween genes, Spook and Phantom in ecdysteroidogenesis in the desert locust, Schistocerca gregaria. J Insect Physiol 57:1240–1248.

Marchal E, Verlinden H, Badisco L, Van Wielendaele P, Vanden Broeck J. 2012. RNAi-mediated knockdown of Shade negatively affects ecdysone-20-hydroxylation in the desert locust, Schistocerca gregaria. J Insect Physiol 58:890–896.

Mashimo Y, Beutel RG, Dallai R, Lee CY, Machida R. 2013. Embryonic development of Zoraptera with special reference to external morphology, and its phylogenetic implications (Insecta). J Morphol 275:295–312.

Miller A. 1940. Embryonic membranes, yolk cells, and morphogenesis of the stonefly Pteronarcys proteus Newman (Plecoptera: Pteronarcidae). Ann Entomol Soc Amer 33:437–477.

Mohamed AA, Zhang L, Dorrah MA, Elmogy M, Yousef HA, Bassal TT, Duvic B. 2016. Molecular characterization of a c-type lysozyme from the desert locust, Schistocerca gregaria (Orthoptera: Acrididae). Dev Comp Immunol 61:60–69.

Mortazavi A, Williams BA, McCue K, Schaeffer L, Wold B. 2008. Mapping and quantifying mammalian transcriptomes by RNA-Seq. Nat Methods 5:621–628.

Miyakawa K. 1979. Embryology of the dobsonfly, Protohermes grandis Thunberg (Megaloptera: Corydalidae), I. Changes in External form of the embryo during development. Kontyû 47: 367–375.

Nijhout HF. 1994. Insect hormones. Princeton: Princeton University Press.

Niwa R, Matsuda T, Yoshiyama T, Namiki T, Mita K, Fujimoto Y, Kataoka H. 2004. CYP306A1, a cytochrome P450 enzyme, is essential for ecdysteroid biosynthesis in the prothoracic glands of Bombyx and Drosophila. J Biol Chem 279:35942–35949.

Niwa R, Niwa YS. 2014. Enzymes for ecdysteroid biosynthesis: their biological functions in insects and beyond. Biosci Biotechnol Biochem 78:1283–1292.

Noh MY, Muthukrishnan S, Kramer KJ, Arakane Y. 2018. A chitinase with two catalytic domains is required for organization of the cuticular extracellular matrix of a beetle. PLoS Genet 14:e1007307.

Norling U. 1982. Structure and ontogeny of the lateral abdominal gills and the caudal gills in Euphaenidae (Odonata: Zygoptera) larvae. Zool Jb Anat 107:343–389.

Novak VJA, Zambre SK. 1974. To the problem of structure and function of pleuropodia in Schistocerca gregaria FORSKÅL embryos. Zool Jb Physiol 78:344–355.

Ou Q, Zeng J, Yamanaka N, Brakken-Thal C, O’Connor MB, King-Jones K. 2016. The Insect Prothoracic Gland as a Model for Steroid Hormone Biosynthesis and Regulation. Cell Rep 16:247–262.

Pagès H, Aboyoun P, Gentleman R and DebRoy S. 2017. Biostrings: Efficient manipulation of biological strings. R package version 2.46.0.

Panfilio KA. 2008. Extraembryonic development in insects and the acrobatics of blastokinesis. Dev Biol 313:471–491.

Pertea G, Huang X, Liang F, Antonescu V, Sultana R, Karamycheva S, Lee Y, White J, Cheung F, Parvizi B, Tsai J, Quackenbush J. 2003. TIGR Gene Indices clustering tools (TGICL): a software system for fast clustering of large EST datasets. Bioinformatics 19:651–652.

Petryk A, Warren JT, Marqués G, Jarcho MP, Gilbert LI, Kahler J, Parvy JP, Li Y, Dauphin-Villemant C, O’Connor MB. 2003. Shade is the Drosophila P450 enzyme that mediates the hydroxylation of ecdysone to the steroid insect molting hormone 20-hydroxyecdysone. Proc Natl Acad Sci USA 100:13773–13778.

Pesch YY, Riedel D, Patil KR, Loch G, Behr M. 2016. Chitinases and Imaginal disc growth factors organize the extracellular matrix formation at barrier tissues in insects. Sci Rep 6:18340.

Prpic NM, Wigand B, Damen WG, Klingler M. 2001. Expression of dachshund in wild-type and Distal-less mutant Tribolium corroborates serial homologies in insect appendages. Dev Genes Evol 211:467–477.

Qu M, Ma L, Chen P, Yang Q. 2014. Proteomic analysis of insect molting fluid with a focus on enzymes involved in chitin degradation. J Proteome Res 13:2931–2940.

R Development Core Team. 2008. R: A language and environment for statistical computing. R Foundation for Statistical Computing, Vienna, Austria. ISBN 3-900051-07-0.

Rau A, Gallopin M, Celeux G, Jaffrezic F. 2013. Data-based filtering for replicated high-throughput transcriptome sequencing experiments. Bioinformatics 29:2146–2152.

Rathke H. 1844. Zur Entwickelungsgeschichte der Maulwurfsgrille (Gryllotalpa vulgaris). Arch Anat Physiol wiss Med 27–37.

Reynolds SE, Samuels R. 1996. Physiology and biochemistry of insect moulting fluid. Adv In Insect Phys 26:157–232.

Rong S, Li DQ, Zhang XY, Li S, Zhu KY, Guo YP, Ma EB, Zhang JZ. 2013. RNA interference to reveal roles of β-N-acetylglucosaminidase gene during molting process in Locusta migratoria. Insect Sci 20:109–119.

Roonwall ML. 1937. Studies on the embryology of the African migratory locust, Locusta migratoria migratorioides Reiche and Frm. (Orthoptera, Acrididae). II. Organogeny. Philos Trans R Soc Lond B Biol Sci 227: 175–244.

Rost MM, Poprawa I, Klag J. 2004. Ultrastructure of the pleuropodium in 8-d-old embryos of Thermobia domestica (Packard) (Insecta, Zygentoma). Ann Entomol Soc Amer 97:541–547.

Schulz MH, Zerbino DR, Vingron M, Birney E. 2012. Oases: robust de novo RNA-seq assembly across the dynamic range of expression levels. Bioinformatics 28:1086–1092.

Shi L, Paskewitz SM. 2004. Identification and molecular characterization of two immune-responsive chitinase-like proteins from Anopheles gambiae. Insect Mol Biol 13:387–398.

Shutts JH. 1952. Some characteristics of the hatching enzyme in the eggs of Melanoplus differentialis (Thomas). Proc S Dak Acad Sci 31:158–163.

Simão FA, Waterhouse RM, Ioannidis P, Kriventseva EV, Zdobnov EM. 2015. BUSCO: assessing genome assembly and annotation completeness with single-copy orthologs. Bioinformatics. 31:3210–3212.

Slifer EH. 1937. The origin and fate of the membranes surrounding the grasshopper egg; together with some experiments on the source of the hatching enzyme. Q J Micr Sci 79:493–506.

Slifer EH. 1938. A cytological study of the pleuropodia of Melanoplus differentialis (Orthoptera, Acrididae) which furnishes new evidence that they produce the hatching enzyme. J Morphol 63:181–206.

Stanley MSM, Grundmann AW. 1970. The embryonic development of Tribolium confusum. Ann Entomol Soc Amer 63:1248–1256.

Stay B. 1977. Fine structure of two types of pleuropodia in Diploptera punctata (Dictyoptera: Blaberiadae) with observations on their permeability. Int J Insect Morphol Embryol 6:67–95.

Sugahara R, Tanaka S and Shiotsuki T. 2017. RNAi-mediated knockdown of SPOOK reduces ecdysteroid titers and causes precocious metamorphosis in the desert locust Schistocerca gregaria. Dev Biol 429:71–80.

Sui Y-P, Liu X-B, Chai L-Q, Wang J-X, Zhao X-F. 2009. Characterization and influences of classical insect hormones on the expression profiles of a molting carboxypeptidase A from the cotton bollworm (Helicoverpa armigera). Insect Mol Biol 18:353–363.

Supek F, Bošnjak M, Škunca N, Šmuc T. 2011. REVIGO Summarizes and Visualizes Long Lists of Gene Ontology Terms. PLoS ONE 6: e21800.

Tanaka M, Kobayashi Y, Ando H. 1985. Embryonic development of the nervous system and other ectodermal derivatives in the primitive moth, Endoclita sinensis (Lepidoptera, Hepialidae). In: Ando H, Miya K, editors. Recent Advances in Insect Embryology in Japan. Tsukuba: Isebu Co. Ltd. p 215–229.

Tanizawa T, Ando H, Tojo K. 2007. Notes on the pleuropodia in the giant water bug Appasus japonicus (Heteroptera, Belostomatidae). Proc Arthropod Embryol Soc Jpn 42:9–11.

Tarazona S, García-Alcalde F, Dopazo J, Ferrer A, Conesa A. 2011. Differential expression in RNA-seq: a matter of depth. Genome Res 21:2213–2223.

Tear G, Akam M, Martinez-Arias A. 1990. Isolation of an abdominal-A gene from the locust Schistocerca gregaria and its expression during early embryogenesis. Development 110:915–925.

Tsutsumi K, Machida R. 2006. Embryonic development of a snakefly, Inocellia japonica Okamoto: an outline (Insecta: Neuroptera, Raphidiodea). Proc Arthropod Embryol Soc Jpn 41: 37–45.

Uchifune T, Machida R. 2005. Embryonic development of Galloisiana yuasai Asahina, with special reference to external morphology (insecta: Grylloblattodea). J Morphol 266:182–207.

Untergasser A, Nijveen H, Rao X, Bisseling T, Geurts R, and Leunissen JMA. 2007. Primer3Plus, an enhanced web interface to Primer3. Nucleic Acids Research 35: W71–W74.

Viscuso R, Sottile L. 2008. Fine structure of pleuropodia in three species of Insecta Orthoptera during embryonic development. Ital J Zool (Modena) 75: 11–19.

Xi Y, Pan PL, Ye YX, Yu B, Xu HJ, Zhang CX. 2015. Chitinase-like gene family in the brown planthopper, Nilaparvata lugens. Insect Mol Biol 24:29–40.

Warren JT, Petryk A, Marques G, Jarcho M, Parvy JP, Dauphin-Villemant C, O’Connor MB, Gilbert LI. 2002. Molecular and biochemical characterization of two P450 enzymes in the ecdysteroidogenic pathway of Drosophila melanogaster. Proc Natl Acad Sci USA 99:11043–11048.

Warren JT, Petryk A, Marqués G, Parvy JP, Shinoda T, Itoyama K, Kobayashi J, Jarcho M, Li Y, O’Connor MB, Dauphin-Villemant C, Gilbert LI. 2004. Phantom encodes the 25-hydroxylase of Drosophila melanogaster and Bombyx mori: a P450 enzyme critical in ecdysone biosynthesis. Insect Biochem Mol Biol 34:991–1010.

Waterhouse RM, Seppey M, Simão FA, Manni M, Ioannidis P, Klioutchnikov G, Kriventseva EV, Zdobnov EM. 2018. BUSCO applications from quality assessments to gene prediction and phylogenomics. Mol Biol Evol 35:543–548.

Wei Z, Yin Y, Zhang B, Wang Z, Peng G, Cao Y, Xia Y. 2007. Cloning of a novel protease required for the molting of Locusta migratoria manilensis. Dev Growth Differ 49:611–621.

Wheeler WMM. 1889. On the appendages of the first abdominal segment of embryo insects. Trans Wis Acad Sci Arts Lett 8:87–140, pls 1-3.

Yan J, Cheng Q, Narashimhan S, Li CB, Aksoy S. 2002. Cloning and functional expression of a fat body-specific chitinase cDNA from the tsetse fly, Glossina morsitans morsitans. Insect Biochem Mol Biol 32:979–989.

Young MD, Wakefield MJ, Smyth GK and Oshlack A. 2010. Gene ontology analysis for RNA-seq: accounting for selection bias. Genome Biol 11:R14.

Zerbino DR, Birney E. 2008. Velvet: algorithms for de novo short read assembly using de Bruijn graphs. Genome Res 18:821–829.

Zhang J, Lu A, Kong L, Zhang Q, Ling E. 2014. Functional analysis of insect molting fluid proteins on the protection and regulation of ecdysis. J Biol Chem 289:35891–35906.

Zhang H, Shinmyo Y, Mito T, Miyawaki K, Sarashina I, Ohuchi H, Noji S. 2005. Expression patterns of the homeotic genes Scr, Antp, Ubx, and abd-A during embryogenesis of the cricket Gryllus bimaculatus. Gene Expr Patterns 5:491–502.

Zhu Q, Arakane Y, Beeman RW, Kramer, KJ, Muthukrishnan S. 2008. Functional specialization among insect chitinase family genes revealed by RNA interference. Proc Natl Acad Sci USA 105:6650–6655.

Zhu KY, Merzendorfer H, Zhang W, Zhang J, Muthukrishnan S. 2016. Biosynthesis, Turnover, and Functions of Chitin in Insects. Annu Rev Entomol 61:177–196.

