## Supplementary_file_1 for "Transcriptomics supports that pleuropodia of insect embryos function in degradation of the serosal cuticle to enable hatching"

**SUPPLEMENTARY INFORMATION - SUMMARY**

The Supplementary information contains:

- 7 Supplementary figures (Supplementary file 1)
- 18 Supplementary tables (Supplementary file 2)
- Supplementary references (Supplementary file 1; for Supplementary figures and Supplementary tables)

**FIGURE S1**

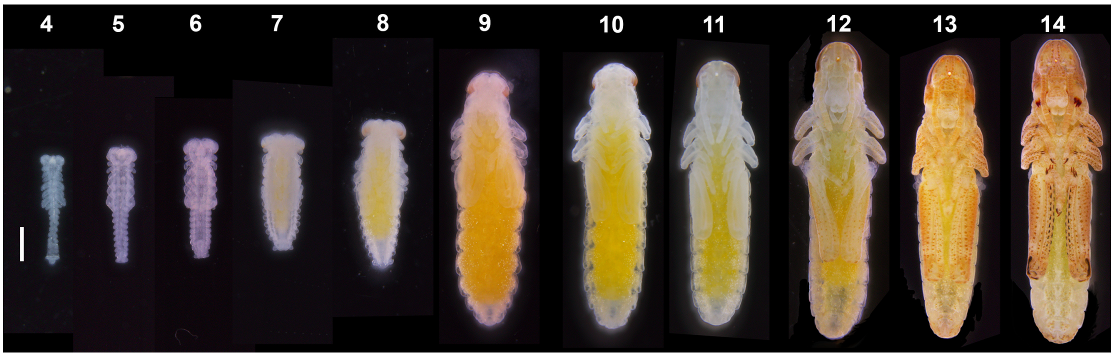


**Figure S1. *Schistocerca* embryonic stages used in this study.** Images of live embryos dissected out of the eggs; imaged under a stereomicroscope. Eggs and embryos of *Schistocerca* typically slightly vary in size. Numbers indicate age in days. Scale bar: 1 mm. Background in photos was cleaned (see Materials and Methods).

**FIGURE S2**

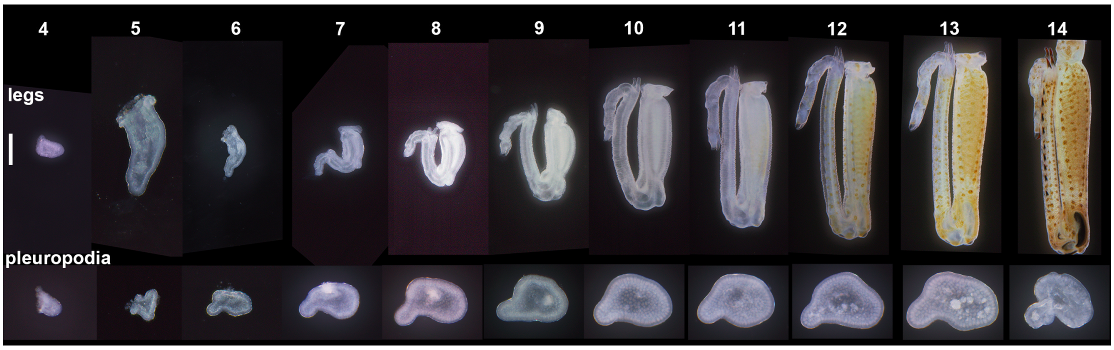


**Figure S2. External features of developing hind legs and pleuropodia.** Compare the sizes of the appendages; imaged under a stereomicroscope. Numbers indicate age in days. Scale bar: 0.2 mm for all pleuropodia and for legs at days 4 and 5; 0.5 mm for legs at days 6-14.

**FIGURE S3**

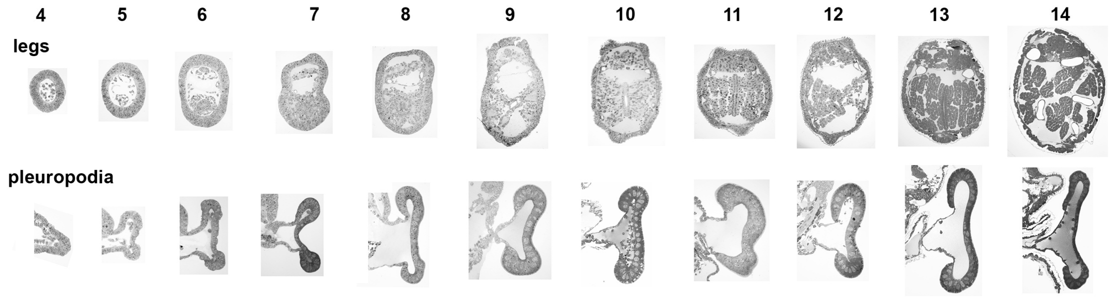


**Figure S3. Cross-sections through developing hind legs and pleuropodia.** Toluidine blue stained semithin sections of appendages embedded in epoxy resin. Numbers indicate age in days.

**FIGURE S4**

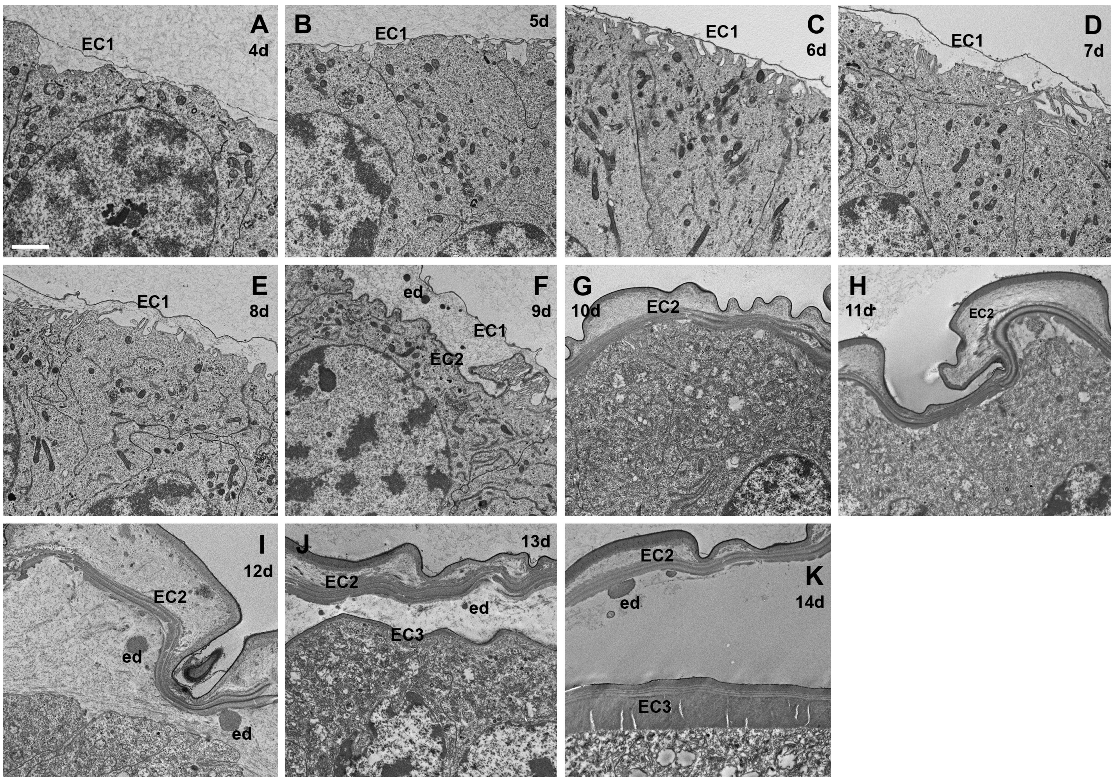


**Figure S4. Ultrastructure of epidermal cells in developing hind legs.** TEM micrographs. Compare with pleuropodia in Figure 3. Note the three different cuticles and appearance of ecdysial droplets (ed) during embryonic moulting.

EC1, EC2, EC3, the first, the second and the third embryonic cuticle, respectively (EC3 becomes the cuticle of the first instar larva). Scale bar: 2 µm.

### FIGURE S5

### (A)

*Sg-nag1*
MSVISTTVLVFALYGIFSCFATQAEEERPVWTWECRESRCEKVAAGEGEAQSLGACRLSCDPWATLWPRPRGGLQRTPGRLLA
LNPYSVSVEAAGRDLQPGVRQLLQEAGRIFHRKVERKARTGAKLRSAGERRSLFVTLTVSDGQTRSFHTDTSEAYSLSISEVTAG
RVNAAVTADTFFGARHALETLYQLIVYDDINKQLLLLSEINLSDSPAFPHRAIALDTARSYFSVASIKRTIDAMAANKLNTFHW
HITDSHSFPFVSETFPKLSQYGAYSPEKVYTPDEIKSVVEYARVRGVRIIPEFDAPAHVGEGWQWVGDNATVCFKADPWSQYCV
EPPCGQLNPTSEKMYQVLAGIYKDMLNVFDSDVFHMGGDEVNMNCWNTSEVITDWMDANGIPRTEEGLHELWDRFQSRAY
SLLAEANGKKELPVILWTSTLTDVAHVDKYLDNKRYIIQIWTRGTDLVIPELIRKGFRVIFSNYDALYFDCGFGAWIGSGNNWCS
PYIGWQKVYDNNVWDLLSAFGIDVGEGSEARKLVLGSEAALWSEQADEFALDGRLWPRAAALAERLWTDPVEGWMSAEHRF
LIQRQRLVDEGIAADTIEPEWCLQNQGHCYA*

*Sg-nag2*
MAPAPPAPHLLALTLLLTLLPSPPVVWANSPRWQWTCDSGLCVRSEAPPEPRLDAELEETVVQRSVHRLRPPWPSHELCRLT
CGPYGALWPRPTGHTLIADALVPFNPATARFDLSAVAGEQGRELVDAASRRWVRDLQHALAASGGHGGGEVAGAAAGAGT
DVLVTVLTRDSPQALSWETDETYTLDVASSGHEVRVTVSAQTVWGALHGLTSLRQLVGCCSEDGAALMVAEARIVDGPVYAH
RGLLLDTARNFLPVETMMATMDAMAASKLNVLHWHATDSQSFPLLLPRVPQLARWGAFSARETYSSQQVSALLGYAHARGI
RLLLELDAPAHSGQGWQWGEAEGLGALALCVGQQPWRRLCIQPPCGQLNPANPRLVGVLADVYRDVVDLWPPGQPLHMGG
DEVSYSCWNSSAEVLEYMSKRRWDRSQDGFLRLWAEFQQAALEALDAARGSSDVPAILWSSHLTRPGNIERFLNSSRYVIETW
VEGGDPLPQQLLALGYRLVVATKDAWYLDHGFWGSTRYHDWKAVYSNRLPGSMAQGVLGGEVASWGELVDDQSLDARLWP
RAAALAERLWSNPGASAREAEPRLHAHRARLVAAGVRPEALAPRYCVLNEGACQ*

*Sg-fdl*
MSRQRLLWRLLGAALALTVAGLAAPPLFRLLVSPHSAANSVAGRRVYSSDPGPWTWSCESGRCVRALWQGGTQVSLDTCQW
TCAGWEAPLWPRPTGALRLANSTAALPEDLDVRLRLSGPQHEDTRGLLAAATERLARHLQLVRPAWAGRVACDAARGATVA
RLTVFVKLDADGSRPTGQLTLDTDESYRLQVRRESQDLQAEIDARSFFGARHALETLSQLAWWDPVSGCVHILDSAIVKDAPKF
RHRGLMVDTARNFIPLEALQRTVDAMASNKLNTLHWHLTDSTSFPYLSRALPTMARYGAYSPEQVYSMEDVSRLAEFARERG
VRLVVELDVPAHAAAGWPTEQVSCSEQRGSAANAPLVQQQQHRQNEDNGLQYRQEERRERRAQHGGEQQPAWWELCGQPP
CGQLPPADEAAFGTLRTLYQELRQASGASDVAHLGGDEVSAECWGGVRGERLWSLWGGFMRRAHRELVAASQGNPPTAVLV
WSSELTAPHNLRRYFDPSTHVVQVWGGSKWNETLPVLLAGFRAVVSHVDAWYLDCGWGDFRSGGPGPCGPVATWQTVYSH
RPWAAFPPGARSRLLGGEACLWSEKVDDQTLDVRLWPRAAALAERLWSDPPAGVHPDLPPPGSPQRDEPTLRRAYQRLSHH
RERLVARGVRAEAMWPRYCHLNPGACF*

*Sg-hex*
MGKKVEVVLCACVCVGLLLTVTAAEPLPRYITEPGPTVKATQGAVWPKPQNEQRFGGSVLIVPGNFTFQVEGPECDILSEAVSR
YEAILKEEAAIKGPRNASEASTQLSALLVRLDGECGDRPVFGMDESYELRINSPDLPGAMLLTSASVWGILRGLETFSQVATRVK
TADALILDNLAIADIPRFSHRGLLLDTSRHFIPVSYIKKTLDAMAYNKMNVFHWHIVDDQSFPYQSAAFPLLSEKGSYDPERFVY
SPADVAEVIEYARVRGIRVVPEFDTPGHTRSWGEAYPDLLTPCYNATGSPDGTYGPIDPTKNFTYEFLQTLFEEIVNVFPDEYFH
LGGDEVGFECWESNQDILDFMSEHNITESKDLESYYIQKIVDIASNLNSKSIVWQEVFDNEVRLSADTVVHIWTGDRNEELDSV
TAAGHYTLLSQCYYLDRFRYFGGDWHKFYNCEPLDFSADNVYQYDLVIGGEAAMWSEFVDESNVESRVWPRASAVAERLWSP
MNVTDIDEAATRIEEHYCRLRRRGINAQPPNGPGYCV*

### (B)

*Sg-cht5-1*
MRTSAAWFLAVAGLCVVFCPPLVSGNVGDRGRVVCYFSNWAIYRPGIGRYGIDDVPASMCTHLVYSFIGVSNVTWGVLVIDPE
NDVENHGFANFTALKSKYPGLKTQLAIGGWAEGGRKYSAMAAVPARRRSLIASVVEYMKRYGFDGFDLDWEYPGAADRGGSF
SDKNHFKCFVQELREAFDAEGQGWEITMAVPLAKFRLQEGYHVPELCELVDAIHVMSYDLRGNWAGFADTHSPLYKRPHDQ
WAYEKLNVHDGLKLWQDMGCPAHKLVVGVPFYGRSFTLSAGNKDYKLGTYINKEAGGGKPGNYTQAKGFLAYYEICLEIQEVG
GWTEKWDEAGKVPYAYKGTQWVGFENPKSVQIKMDFIKAKGYGGAMTWAIDMDDFRGVCGPKDALISVMYNNMKDYIVPD
IQYSTTKRPDWDRPPPCDGKKPGAAPASTTTRRPTAAPTQSTTRRPAPTTTAAPSSSSSTTTTRRTTTASRPSTQPPPPPAAPD
DNELPPAAIDCSDGDFVPHHDCSKYYRCVYGKPVEFSCYEGTVWNPQLRVCDRPNDVHRTDCSMAKLHS*

*Sg-cht5-2*
MRAATQVGLLLAVALALAAASDEDTTPLDSSTGSPTNSVDEESSSSENAAVLSGGQRRGRVTCYFESWAVYRKRLRYGIEDIPG
DMCTHIIYSFVGLNNVTWELQVLDEKLDVQDGGFENFTALRQEFPGVRLQVALGGWAEGGHNYSAMVGDPARRASLVRSAVA
FLHRYGFDGFDVDWEYPGNAPRGGVPEDKDDFLCFMQELRVAFDAEGLGWELTMAVPLTEDKLRDGFHVPQLCSIVDAVHV
MAYDLRGEWDHFADVHSPLYRRPHDTGAYAKINTHDGLLLWEQLGGSSWGCHSTATPTNCVPTSPTTLPVLASFRAPEMTSA
E*
(based on alignment with homologous sequences this transcript might be misassembled and the amino acid sequenced prematurely terminated by introduction of a stop codon)

*Sg-cht10-1*
MWRPVALSLWLLLATSRGLHVPPADEPSFVRDAVEAPPGQSLALRRSATASRPRLPAFGTRQLPLRQAVESPPMAARLRSSER
LPLRDAVEHVPYEALPGAPTASEAFSLWRGFGDWLPENLPSTRQFNHSFAWWHDAIIAKLSLGGPRTKPPSLQAPSTHTSGIR
QFKVVCFVEGWAGYRRDPMRFTTADIDPFACTHIIYAFAVMDPHDLHIKPQDEQYDIIQGGYRSIVGLKRQNPQLKVMISVGG
WPEERRKFAEMTASASTRREFIRSVLHFIDEYGFDGIDLDWEYPGAADMGGSAREKEHFSLLVEELAEAFAPRGSVLSASVSPS
RFRVEDGYDVPRLARRLDFLNLMAFDLLTEQDAAADHHAPLTQRKHDYGLAVFYNVDYAVRYWLRKGARRDQLVVGIPFHG
HSFTLQDEAKNSPGAPVKGLGKEGPYTQEKGFLAYFEILQLLEEGHWMKATDDVGSPYMVKGNQWIGYEDQRSIATKVMYIK

KNLLGGAMVWALDLDDFEGAYGQKWPLLSVVKKGLLETTPQSDQQQASQEPTHVTPPIAGVPVSVDSSQYNCSGRGYVRDSA
SCQIYHRCEWGMKHTYICPEGLHYDSRTQLCDWPQIANCPMDNSSQRIEQENQSEVACNEEGLMEDPKDCNRYYMCHKGVA
QHYSCMLGQYFNVQKGICEYGSCMPKAPQDNIPSSQTRNLVGEDHYKVVCYYASWAWYRKEGGKFVPEHIDPTLCTHIVYAYA
SLDPNTLTMKYFDERADKENNFYERLTELPKKSGHHQQHASDVTVMIGLGGWTDSAGDKYSRLVSEGSARRRFVSKAVEFLH
RHQFGGLHLDWDYPRCWQSNCGRGPTSDKPNFTKLVQELRQAFKKQNPPLALAISISGYHEVIDEAYDLAELGRNTDFMSVM
TYDYHGSWEKSTGHVSPLYHRNGDIFPMYNTNDTMEYLVNKGAPRDKLLVGIPFYGQSYTLENPSNHDIGAPATGPGLAGEFT
MQPGMLAYYEICDRVRNNFWKIGRDRFGATGPFAYAGNQWVSFEDTKSVKEKAKYIKNMGYGGAMTFTLDLDDFENRCCRG
AFPLLRSINRVFGRIPDSAEPSGDDCTRPPPPVTPPPPTYTTGVDSGDHRPTTPISTTHQHPTSPKPSTTEYPWW

*Sg-cht10-2*
PSTTTSTTTSTTTTTTTTTTSTTTTPRPTTRPTTMSTTEYPWWTPSTTSTTRKPPTTRPTTTSTTEEYPWWTPPSTTKKPTTS
STTEQPWWTPSTTSTTSTAAPTTTMTTTEKPWWSTTPQKPLPPDSGPCEAGVYYPDPTNCNAYYRCVLGELRKEFCAGGLH
WNPDKKVCDWPSESKCDTKEPSETTVGSTTSSTTENPWWTPSKPSETQATTTTTEVPWWSTTRPPRPPTTEGNSEWVTTSR
PTTTQQPSEEVSECMNGQYYPVAGSCKSFYICVNGRLIKQTCAPGLVWNQDQTMCDWGFNVKCADDSEREAVHKAQPDDPC
NQGALNPYPGDCTRYLYCQWGRYHEADCAAGLHWNEMEKICDWPENAKCTDMESGSEAPAASSQKPVTEMSTSWTTAAPT
TKPPWTWATTTTVKPVTTTSTRAPPAQGPPISGYFKVVCYFTNWAWYRRGLGKYVPEDIDANLCTHIVYGFAVLDYENLIIKA
HDSWADFDNKFYERVVAYKKKGLKVSLAIGGWNDSAGDKYSRLVNSPSARRRFIKHVLEFLEKYGFDGLDLDWEYPVCWQVD
CAKGPASDKSSFAALVKELRQAFEPKGLLLSSAVSPSKTVIDAGYDVKTLAENLDWIAVMTYDFHGQWDKKTGHVAPLYFHPD
DDFYFFNANFSINYWISEGAPRRKIVMGMPLYGQSFQLEKASTNGLNARSTGPGQAGEFTRAAGFLAYYEICDRIKNKGWTVV
QDPERRMGPYAFKGNQWVSFDDVAMIQQKSEYIRKMGLGGGMIWALDLDDFRNRCGGGTHPLLNTIRTVLAAPPGGDGATE
MPPSWSTPGGGQPTMSTEEWMSSTSISSTEITDSGHHSTQDSGGEVTSVSPAITTTNRPAHPGTSSSPPPPPSQGEFKVVCYFT
NWAWYRQGVGKYLPNEIDPDLCTHIVYGFAVLNGDRLTIKPHDTWADYDNKFYEKVTEYKKKGIKVLVAIGGWNDSAGDKYS
RLVNSPGARRRFIEDVIDFIEQNNFDGLDLDWEYPKCWQVDCKKGPDSDKEAFAAFVRELRAAFNPKGLLLTAAVSPSKAVVD
AGYDVPTLSQNLDWIAVMTYDFHGQWDKITGHVAPMYTHPEDVDVTFNANFSIHYWIQKGASPKKIVMGMPMYGQSFSLAD
NSDHGLNAPTYGGGEAGESTRARGFLSYYEICTNIQKKGWRVVKDPEGRMGPYAYLRDQWVSFDDTSMIRYKSNFIRRMGLG
GGMIWALDLDDFRNVCSCEKYPLLKTINRVLRGYPGPGPNCDIEATEKPGSEETDNRIHPTIPPTSSTNNWNVISGGGGIVPKDP
TCGNRLFAPHDKDCNKYYLCQYGDFMEQSCPQGLYWNKDHCDWPSNTDCSKEDSSVINPAPIASTQEPEMSSTTENIHMSES
TVTTSIRPSEPGTSTVMTPSGDYMVVCYFTNWAWYRQGLGKYLPSDIDTSLCTHIAYGFAVLDGNSLTIKPHDSWADLDNEFY
TKVSGLKKKGIKVLLAIGGWNDSLGDKYSRLANNPSARRKFVEHVVKFIEKYGFEGLDLDWEYPKCWQVDCNAGPDSDKQGF
ADLVKELSMAFKPRGLLLSSAVSPSKVVIDSGYDVPVLSQYFDYISVMTYDFHGHWDKQTGHVAPLYYYPGDTYDYFNANFTM
HYWIEKGADRKKLIMGMPMYGQSFSLADAKNHGLNAKSYGPGEAGEFTRAGGFMAYYEICYNVKSKGWTTVRDPEGRIGPYA
YRGNQWVSYDDVSDIRRKTQFIKELGLGGGMIWALDLDDFRNRCGCGTYPLLRTINSELRGLTANTHDCT*

*Sg-cht7-1*
MIAPRCVWRAALWCVVIILLADLVYSASSTGRRRLRRPGGSSSSSTTSSSSSTSTKVRTRDQETSASVNRFRVRNRLTPPGANRK
SGSGSAVAAASDKSGGYKVVCYYTNWSQYRTAHGKFLPEDITPDLCTHIIYAFGWLKKGKLTSFEGNDETKDGKVGLYERVMA
LKKANPKLKVLLALGGWSFGTQKFKAMSETRYTRQTFIYSAIPYLRKHDFDGLDMDWEYPKGTDDKKNFVLLLKELREAFEA
EAQEVKQSRLLLSAAVPVGPDNVRGGYDVPAVASYLDFINLMAYDFHGKWERETGHNAPLYAPSSDSEWRKQLSVDHAATM
WVKLGAPKEKLVIGMPTYGRTFTLSNPSNFKVNAPASGGGKAGDFTKEGGFLAYYEVCDMLKKGATYIWDDEMKVPYAVMG
DQWVGFDDERSIRHKMKWLKEGGYGGAMVWTVDMDDFTGTVCGGGVKYPLIGAIREELRGVSRGPNAKDVDWSKVARTVS
LEATTKPAPIKIDVSEVLNRVRKPTKQAPADLSNEVIDLNSRPAQVFCYMTSWSGKRPGAGKFSPEDVDPSLCTHVVFAFATLK
DHKLAPANDKDDGLYERVIALREKNPQLKVLLAIGGWAFGSTPFKELTSNVFRMNQFVYDAIELLRDFKFDGLDVDWEYPRG
ADDRAAYVSLLKELRMAFEGEAKTAEQPRLLLSAAVPASFEAIAAGYDVPEISKYLDFINVMTYDFHGQWERQVGHNSPLYPLE
SATSYQKKLTVDFSAREWVKQGAPKEKLLIGMPTYGRSFTLVDTSKFDIGAPASGGGAAGRYTAEAGFMAYYEVCDFLHHDNT
TLVWDNEQQVPFAYRGDQWVGFDDERSLKTKMGWLKELGFGGIMVWSVDMDDFRGQCGAGKYPLLTSMRQELRDYRVQLE
YDGPYESRGPLGAYTTKDPTSVSCEEEDGHISYHPDKADCTMYYMCEGERKHHMPCPSNLVFNPNENVCDWPENVEGCMHH
TQAPPAARRR*

*Sg-cht7-2*
MTWPPPPLLLSLLVLLATSASARFVSTHDVTPCAVEALAPSDKALLCYYEGRLSVYQLDPCLCTHIVFKDAAVVSDNFGLKIVSD
VSGASLLRARSPSLRTVLGLRLSGAVARAALASPSRRLALARDAARRLYAHHLDGIELSVDDDEAASAAAADAAPAATARQGLV
ALLKALRTALDSHGREKRDYLVSEQVFDDFTTQEYEPTWSDGSSRKSRRRATTTTTTTSTTESPEETAARYLELERDAQNAQL
LLSLPTKPETIAKRYDVKNITRYVDYVVLRTQAMTDDSERGLVYHPSRLMGLDDMLNADAVVDLVTSLGASPAQLVITLPGQA
TAFELRREDRTEPRSPASGAPRTISQPELCRALSRGNWTLERDEDQTAPYAYSGRRWIAFDDALSASIKGKYAVVRGLAGTAVD
AADALDWQGTCGAPASQLRALHSALAQLRRSSRGALLHGLE

*Sg-cht7-3*
DKGMPKNKIIVGIPTYGHSFRLINAENHGWSAPASGYGKIGSKGFVSYPEVCQFLHSTGSKYIFDKNFEVPYAYQGLEWISYDDE
CSVMYKAKYIASSSYGGAMVFSLNVDDHQGVCAGTTFLLTTQIRNILGVSWQ*

*Sg-cht2*
MQQLAPLAFVLAFLAAAFAASPLGHNKAVVCYVSSWAVYRPGNGVFTVSDINPNICSHLVYAFAGLNATDNTIITLDKYNDLEE
DYGKGNYKKITGLKNQYPHLKVSIAIGGWNEGSANYSHMASTPTTRQQFIRSVVNFLRKYNFDGLDLDWEYPTQRGGVPSDRE
NFVALVRELRQEFDKNGWLLTAALGASTAVIEKAYDVPMLGKYLDYMHIMCYDYHGTWDKMTGANAPLYGSSPSDTLSVDN
SIRYYLKLGAPAKKLLMGVPLYGRTFMSDANANMGGLGAPAEEKSFQGPYTKEDGYMGYNEICLELKTNSSMWTIMWDDKSS
TPYAVSTNKVIVYDNAKSLTEKVNLAMKLELGGIMVWPLDTDDFRGECSEGIYPLMHTINKAIVQSSQQKSDSSGMKVPDSTA
AASCGCASLIFLSFLYLFQL*

*Sg-cht6-1*
VCYYTNWSVYRPGTAKFTPQNINPYLCTHLIYAFGGLSRENGLRPFDKYQDIEQGGYAKFTGLKTYNKDLKTMLAIGGWNEGS
TRFSPLVADAERRKEFVKNVLRFLRQNHFDGLDLDWEYPAFRDGGKSRDRDNYALLVKELREEFDRESEKTGRPRLLLTMAV
PAGIEYIDKGFDIASMNKHLDFMNILSYDYHSAFEPAVNHHSPLYSMEEDDEYNFDAQLTIDHTVNHYMKSGADRNKLVLGIP
TYGRSYTLFNPLATELGSPADGPGEQGDSTREKGYLAYYEICENLQSDDWKVVQPNPSAMGPYAYKGNQWVSYDDMDIIKKK
AQYVNDNGLGGIMFWAIDNDDFRGKCHGRPYPLIEAGKEAMLKGVKRSNNEIETTPVQNNRQSSRKRNRNRSKGNARGRTRT

TASTSTVVTTTTTTTTTTAAPLITPSYTTPEPPTTPDPGSDFKCKDEGFFPHPRDCKKYFWCLDSGPSNLGIVAHQFTCPSGLFF
NKAADSCDYARNVVCNKKSKSQGGSSSTLPPIKAATSSTTRFSTSPSTKLTTKLTTTTTTEPPPVLDDDDDDD

*Sg-cht6-2*
MNIRVKQPVIIGNCYRGQPNRLWEVFILKWFLVAVACLIAAGAVTVYLAHYFMKTRYTSTNVTGVTGQHSDLNTYKGQLQDM
GDGYSLFKQEDMTQICKTDELTGSQQMRKQSTKLVCYYTFPGPGGLVPDKIDPFLCTHINIAAVGINNSKLEPLCEERKEVIKSL
VGLKTRNKNLKVILSVIGMPGGFGDMVSKSSSRRMFIKDL

*Sg-cht8-1*
MSHFWLRLAVILGVSLSICGAEDKKVVCYHGSWSAYRNGNGRFEIEYIQPELCTHLIYTFVGITSAGEVRILDEWLDLPSGKNAY
NRFNALKSSNTKTLVAIGGWNEGSATYSAVMNDASLRAKFVQNVVNFVKTYGFDGFDLDWEYPANRGGSPGDLTAFVELIKE
LRTEFDKYGYLLTAAVGVGRYLIGTAYDVPQISKYLDFINLMTYDLHGSWDGKTGQNAPLYASSADKTEAERQLNVDSSVRYW
IQNGADPSKLVLGMGTYGRTFTLSSAANTGVGAPATAPGTNGPYTMESGMMGYNEICEKINAGGWTVVWDEEQKVPYAVNG
NQWIGYDNEESIRLKSQYVLDMGLAGGMIWSLETDDFKGLCGSKTYPLLSTINEVLRGITSTNGSSSSSSSSSSSSSSSSSSSSSSS
SSSSNTAASASSSSGVCSSAGYVRDPSDCGVFYLCTASGSGYTASKFTCPGDLVFDESSSACNYKSLVAC*

*Sg-cht8-2*
MSPFLSGLLLLLGVLNICGADEKKVVCYHGSWSAYRNGNGRFEIEYIRPELCTHMIYSFVGITSAGEVRILDEWLDLASGKNAYN
RFNKLKSSNTKTLVAIGGWNEGSATYSAVMNNAALRQKFVQNVVNFVKTYGFDGFDLDWEYPANRGGSPGDLRAYVELLKEL
RAEFDKHGFILSAAVGVGRYLIGSAYDVPQLSKYLDFINL

*Sg-cht8-3*
ESGMMGYNEICEKIKAGGWKVTWDDEQKVPYAVSGNQWVGYDNEESIKLKSQYVLDMGLGGGMIWSLETDDFKGVCGAGTF
PLLSAINQVLRGAAATSSAGSSTSGSSSGSSGSSSSSSASSGGSSSGTSSGSSTTSSGASADSSGSSASSGSSSSGSSATSVSSGSNSSG
VCNSAGYARDPSDCGV

*Sg-idgf-1*
MAELPLLLLLLAAAATCWTSAAALGATRVVCYLDGGALRRPEPHRMLVSEIEPSLTYCTHLIYGYATIDTDSYKAVPRHEGEGT
NYTSVVALKRRFPALNVLLSIGGGSADSGQREKYLHLLESDEHRRTFVKSAKDLLKQYHFDGIDIAWEFPMNKEKKERSTLGSF
WHGFKKVIGLAHSHKDEKADEHRREFSSLIQELKTSLKTENALLTLSVIPYINHTLYYDCSALSPHVDHLHLLAYDYHTPQRTP
NTADYPAPLYVAGKRDPDLTADGNVRWFLERGFPSRKIILGIPTFARTWKLDDDSRVSGVPPIEADGAGDTDNIANTAGIMAF
QTVCMLLPNAGNAGYKTTLSRVTDPTDRLGSYGFRLPSGEVTGLWVGYEDPDVAQYKAAYAKIKSLGGIAFSDLSLDDYHGICT
GDKYPIVRAGTLKLRYK*

*Sg-idgf-2*
MQSFARLLLLSACCWSAALAATTKVVCYFNTSALKRPESSRMLLSQIEPSFSYCTHLVVGYATINTETYKAVPPSEDEHTTYTNI
VALKRRFPSLKILLSIGGGAADTDTREKYFELLESDEHRTTFVSSAKSLLKQHGFDGIDIAWEFPKNKAKKDRGTFGSIWHGIKK
AVGAAHSHTDEKADEHKSQFSALIRELRTSLRNENALLTLSVIPYINQSLYYDPTALNQQIDELHVLAFDYRNPERDSQGGRLPC
AALPSRAEGLRPLGRREHPLVPRELIPS*

*Sg-idgf-3*
TDSYKAVPRYQDDTTKYTSLVALKERFPSLKVLLSIGGGGADADQRKKYLELLESDEHRRTFVDSVKELLQQNRFDGIDIAWEFPFSK
EKKDRGNVWHGVKKVLGYAHSHRDENPDEHRRQFSALIRELKSSLKTQNALLTLSVIPYINHSLYYDCASLSPEIDQLHLLAYDYHSP
TRTPKKADYPAPLYRAGERSADLTVDGNVRWFLEKGFPSRKIILGIPTFARTWKLTKDSRITGVPPIDADGPGVAGSIANISGLLAYQT
VCTLLPNDANAAYRTTLRRVTDPTDRLGSYGFRLPTREVSGLWVGYENQHSAEYKAAYARKKSLGGIAFSDLSLDDYNGVCTGEKFP
IVRAGTLKLLSTSV*

**Figure S5. Amino acid sequences and conserved domains of *Schistocerca* chitin degrading enzymes.** (A) NAGs, (B) CHTs. Signal peptide and transmembrane region identified by Phobius (http://phobius.binf.ku.dk/index.html) and conserved domains identified by SMART (http://smart.embl-heidelberg.de/) are underlined and coloured. In (A) and (B): signal peptide: magenta, transmembrane region: dark blue. In (A): Glycohydro 20b2 domain (N-terminal domain of the eukaryotic beta-hexosaminidases): light green, Glyco hydro 20 domain (glycoside hydrolase family 20 catalytic domain): grey. In (B): Glyco 18 domain (catalytic domain): light blue, Chitin-binding domain type 2 (ChBD2): green; catalytically critical consensus sequence in the Glyco 18 domain, FDG(L/F)DLDWE(Y/F)P, is highlighted in yellow and amino acid changes from the consensus are coloured in orange.

**FIGURE S6**

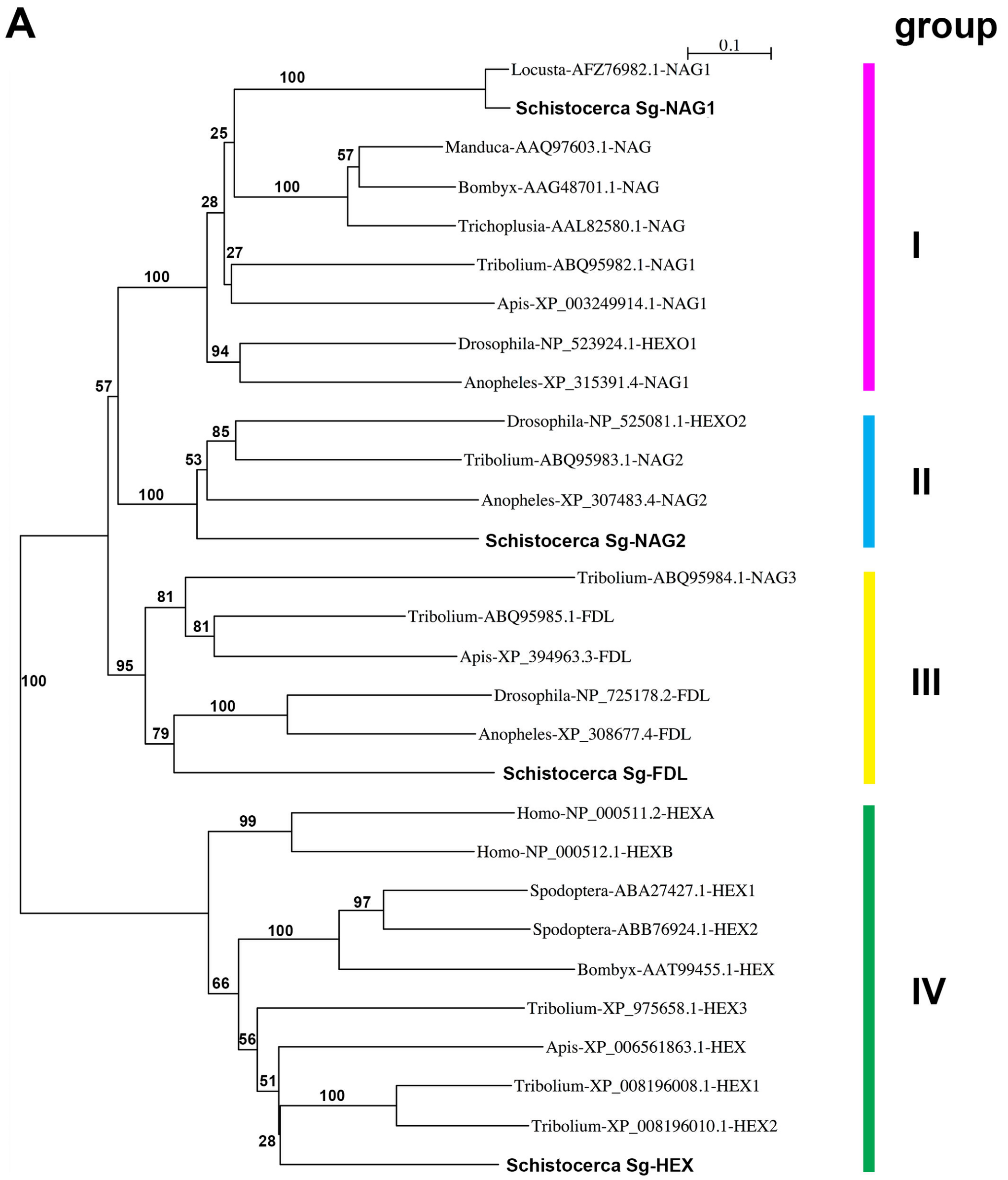
A
group
0.1
Locusta-AFZ76982.1-NAG1
Schistocerca Sg-NAG1
Manduca-AAQ97603.1-NAG
Bombyx-AAG48701.1-NAG
Trichoplusia-AAL82580.1-NAG
Tribolium-ABQ95982.1-NAG1
Apis-XP_003249914.1-NAG1
Drosophila-NP_523924.1-HEXO1
Anopheles-XP_315391.4-NAG1
Drosophila-NP_525081.1-HEXO2
Tribolium-ABQ95983.1-NAG2
Anopheles-XP_307483.4-NAG2
Schistocerca Sg-NAG2
Tribolium-ABQ95984.1-NAG3
Tribolium-ABQ95985.1-FDL
Apis-XP_394963.3-FDL
Drosophila-NP_725178.2-FDL
Anopheles-XP_308677.4-FDL
Schistocerca Sg-FDL
Homo-NP_000511.2-HEXA
Homo-NP_000512.1-HEXB
Spodoptera-ABA27427.1-HEX1
Spodoptera-ABB76924.1-HEX2
Bombyx-AAT99455.1-HEX
Tribolium-XP_975658.1-HEX3
Apis-XP_006561863.1-HEX
Tribolium-XP_008196008.1-HEX1
Tribolium-XP_008196010.1-HEX2
Schistocerca Sg-HEX
I
II
III
IV

B

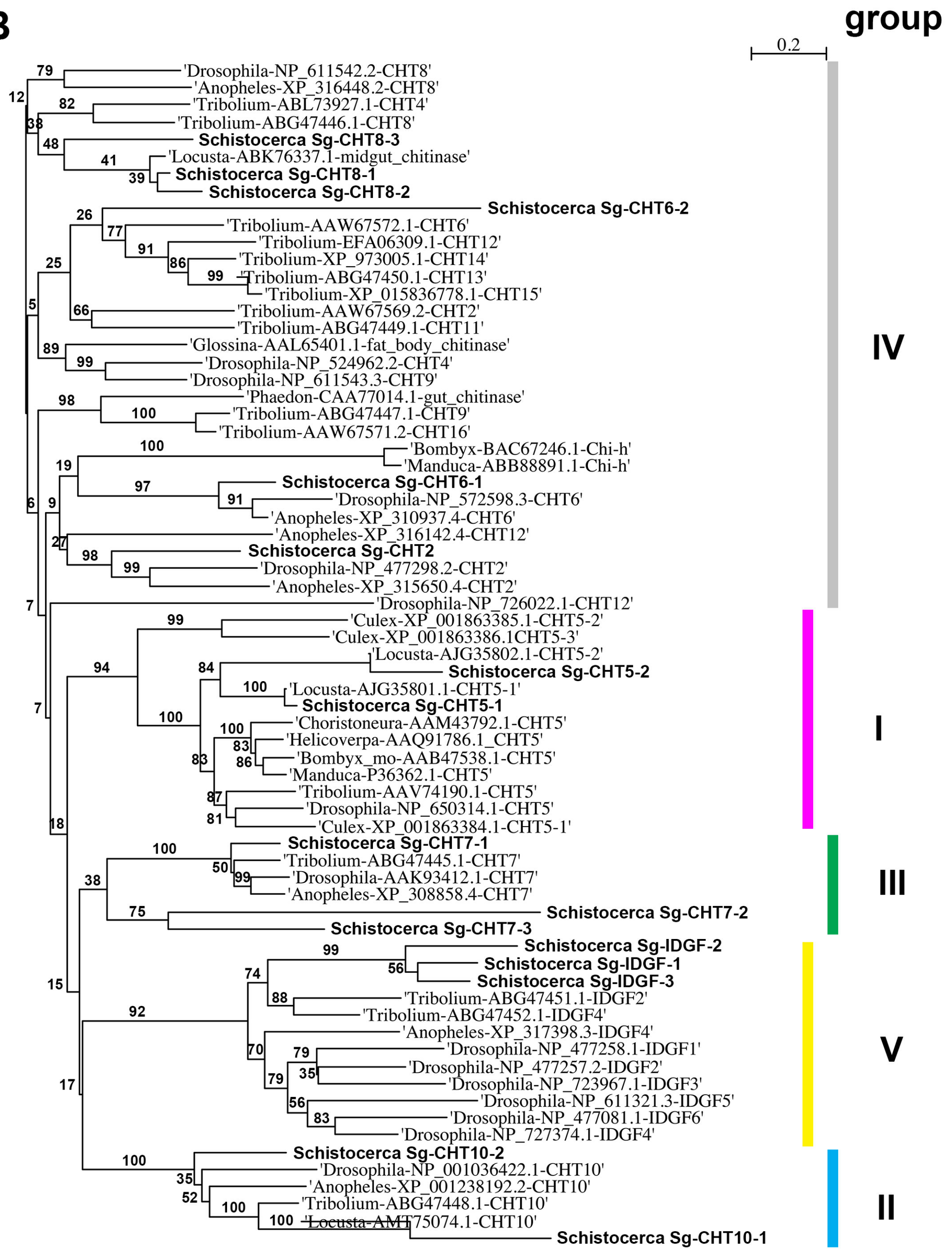
group
0.2
'Drosophila-NP_611542.2-CHT8'
'Anopheles-XP_316448.2-CHT8'
'Tribolium-ABL73927.1-CHT4'
'Tribolium-ABG47446.1-CHT8'
Schistocerca Sg-CHT8-3
'Locusta-ABK76337.1-midgut_chitinase'
Schistocerca Sg-CHT8-1
Schistocerca Sg-CHT8-2
Schistocerca Sg-CHT6-2
'Tribolium-AAW67572.1-CHT6'
'Tribolium-EFA06309.1-CHT12'
'Tribolium-XP_973005.1-CHT14'
'Tribolium-ABG47450.1-CHT13'
'Tribolium-XP_015836778.1-CHT15'
'Tribolium-AAW67569.2-CHT2'
'Tribolium-ABG47449.1-CHT11'
'Glossina-AAL65401.1-fat_body_chitinase'
'Drosophila-NP_524962.2-CHT4'
'Drosophila-NP_611543.3-CHT9'
'Phaedon-CAA77014.1-gut_chitinase'
'Tribolium-ABG47447.1-CHT9'
'Tribolium-AAW67571.2-CHT16'
'Bombyx-BAC67246.1-Chi-h'
'Manduca-ABB88891.1-Chi-h'
Schistocerca Sg-CHT6-1
'Drosophila-NP_572598.3-CHT6'
'Anopheles-XP_310937.4-CHT6'
'Anopheles-XP_316142.4-CHT12'
Schistocerca Sg-CHT2
'Drosophila-NP_477298.2-CHT2'
'Anopheles-XP_315650.4-CHT2'
'Drosophila-NP_726022.1-CHT12'
'Culex-XP_001863385.1-CHT5-2'
'Culex-XP_001863386.1CHT5-3'
'Locusta-AJG35802.1-CHT5-2'
Schistocerca Sg-CHT5-2
'Locusta-AJG35801.1-CHT5-1'
Schistocerca Sg-CHT5-1
'Choristoneura-AAM43792.1-CHT5'
'Helicoverpa-AAQ91786.1_CHT5'
'Bombyx_mo-AAB47538.1-CHT5'
'Manduca-P36362.1-CHT5'
'Tribolium-AAV74190.1-CHT5'
'Drosophila-NP_650314.1-CHT5'
'Culex-XP_001863384.1-CHT5-1'
Schistocerca Sg-CHT7-1
'Tribolium-ABG47445.1-CHT7'
'Drosophila-AAK93412.1-CHT7'
'Anopheles-XP_308858.4-CHT7'
Schistocerca Sg-CHT7-2
Schistocerca Sg-CHT7-3
Schistocerca Sg-IDGF-2
Schistocerca Sg-IDGF-1
Schistocerca Sg-IDGF-3
'Tribolium-ABG47451.1-IDGF2'
'Tribolium-ABG47452.1-IDGF4'
'Anopheles-XP_317398.3-IDGF4'
'Drosophila-NP_477258.1-IDGF1'
'Drosophila-NP_477257.2-IDGF2'
'Drosophila-NP_723967.1-IDGF3'
'Drosophila-NP_611321.3-IDGF5'
'Drosophila-NP_477081.1-IDGF6'
'Drosophila-NP_727374.1-IDGF4'
Schistocerca Sg-CHT10-2
'Drosophila-NP_001036422.1-CHT10'
'Anopheles-XP_001238192.2-CHT10'
'Tribolium-ABG47448.1-CHT10'
'Locusta-AMT75074.1-CHT10'
Schistocerca Sg-CHT10-1
IV
I
III
V
II

244

**Figure S6. Phylogenetic trees of chitin degrading enzymes in *Schistocerca* and other insects.** (A) NAGs, (B) CHTs. *Schistocerca* sequences are in bold. Amino acid sequences were extracted from NCBI GenBank. The numbers above the branches are bootstrap support. The markers show a branch length. Both trees are unrooted. The tree in (A) was prepared using the SeaView software (version 4.6.1; Gouy et al, 2010; http://doua.prabi.fr/software/seaview): alignment with default parameters, tree using the Neighbor Joining method, Poisson distribution, 5000 bootstrap replicates. The tree in (B) was prepared using the CLC Sequence Viewer (version 7.8.1; https://www.qiagenbioinformatics.com/products/clc-sequence-viewer/): alignment with default parameters except gap open cost 3.0 and gap extension cost 3.0, tree using Neighbor Joining method, Kimura model, 1000 bootstrap replicates.

**FIGURE S7**

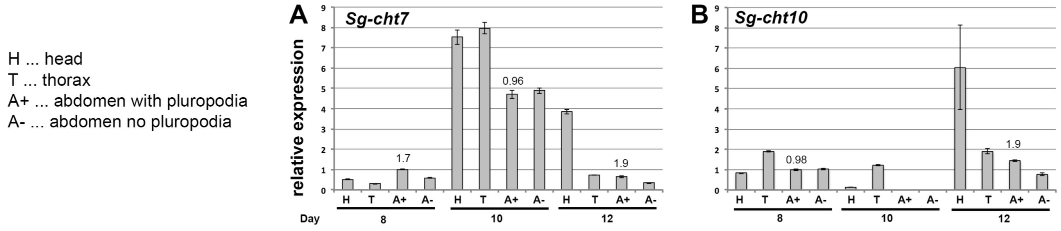


**Figure S7. Real-time RT-PCR expression analysis of *Sg-cht7-1* and *Sg-cht10-1* on cDNA from parts of *Schistocerca* embryos.** cDNA was prepared from mRNAs isolated from parts of embryos at the age of 8, 10 and 12 days: H, head; T, thorax; A+, abdomen with pleuropodia; A-, abdomen without pleuropodia. Analysis of 3-4 technical replicates is shown. Expression in A+8 (abdomen with pleuropodia when they first become differentiated) was set as 1. Numbers above A+ expression is fold change from A- of the same age.

### SUPPLEMENTARY REFERENCES


Angelini DR, Liu PZ, Hughes CL, Kaufman TC. 2005. Hox gene function and interaction  in the milkweed bug Oncopeltus fasciatus (Hemiptera). Dev Biol 287:440-455.

Beermann A, Jay DG, Beeman RW, Hülskamp M, Tautz D, Jürgens G. 2001. The Short antennae gene of Tribolium is required for limb development and encodes the orthologue of the Drosophila Distal-less protein. Development 128:287-297.

Bennett RL, Brown SJ, Denell RE. 1999. Molecular and genetic analysis of the Tribolium Ultrabithorax ortholog, Ultrathorax. Dev Genes Evol 209:608-619.

Gouy M, Guindon S, Gascuel O. 2010. SeaView version 4: a multiplatform graphical user interface for sequence alignment and phylogenetic tree building. Mol Biol Evol 27:221-224.

Hughes CL, Kaufman TC. 2002. Hox genes and the evolution of the arthropod body plan. Evol Dev 4:459-499.

Kelsh R, Weinzierl RO, White RA, Akam M. 1994. Homeotic gene expression in the locust Schistocerca: an antibody that detects conserved epitopes in Ultrabithorax and abdominal-A proteins. Dev Genet 15:19-31.

Liu HW, Wang LL, Tang X, Dong ZM, Guo PC, Zhao DC, Xia QY, Zhao P. 2018. Proteomic analysis of Bombyx mori molting fluid: Insights into the molting process. J Proteomics 173:115-125.

Prpic NM, Wigand B, Damen WG, Klingler M. 2001. Expression of dachshund in wild-type and Distal-less mutant Tribolium corroborates serial homologies in insect appendages. Dev Genes Evol 211:467-477.

Tear G, Akam M, Martinez-Arias A. 1990. Isolation of an abdominal-A gene from the locust Schistocerca gregaria and its expression during early embryogenesis. Development 110:915-925.

Wei Z, Yin Y, Zhang B, Wang Z, Peng G, Cao Y, Xia Y. 2007. Cloning of a novel protease required for the molting of Locusta migratoria manilensis. Dev Growth Differ 49:611-621.

Zhang J, Lu A, Kong L, Zhang Q, Ling E. 2014. Functional analysis of insect molting fluid proteins on the protection and regulation of ecdysis. J Biol Chem 289:35891-35906.

Zhang H, Shinmyo Y, Mito T, Miyawaki K, Sarashina I, Ohuchi H, Noji S. 2005. Expression patterns of the homeotic genes Scr, Antp, Ubx, and abd-A during embryogenesis of the cricket Gryllus bimaculatus. Gene Expr Patterns 5:491-502.
